# CSDE1 is a Post-Transcriptional Regulator of the LDL Receptor

**DOI:** 10.1101/2020.08.03.235028

**Authors:** Geoffrey A. Smith, Arun Padmanabhan, Bryan H. Lau, Akhil Pampana, Li Li, Y. Clara Lee, Angelo Pelonero, Tomohiro Nishino, Nandhini Sadagopan, Rajan Jain, Pradeep Natarajan, Roland S. Wu, Brian L. Black, Deepak Srivastava, Kevan M. Shokat, John S. Chorba

## Abstract

The low-density lipoprotein receptor (LDLR) controls cellular delivery of cholesterol and clears LDL from the bloodstream, protecting against atherosclerotic heart disease, the leading cause of death in the United States. We therefore sought to identify regulators of the LDLR beyond the targets of current clinical therapies and known causes of familial hypercholesterolemia. We show that Cold Shock Domain-Containing Protein E1 (CSDE1) enhances hepatic *LDLR* mRNA decay via its 3’ untranslated region to regulate atherogenic lipoproteins *in vivo*. Using parallel phenotypic genome-wide CRISPR interference screens, we found 40 specific regulators of the LDLR left unidentified by observational human genetics. Among these, we show that CSDE1 regulates the LDLR at least as strongly as the mechanistically distinct pathways exploited by the best available clinical therapies: statins and PCSK9 inhibitors. Additionally, we show that hepatic gene silencing of *Csde1* treats diet-induced dyslipidemia in mice better than that of *Pcsk9*. Our results reveal the therapeutic potential of manipulating a newly identified key factor in the post-transcriptional regulation of the *LDLR* mRNA for the prevention of cardiovascular disease. We anticipate that our approach of modelling a clinically relevant phenotype in a forward genetic screen, followed by mechanistic pharmacologic dissection and *in vivo* validation, will serve as a generalizable template for the identification of therapeutic targets in other human disease states.

**One Sentence Summary:** A genome-wide CRISPR screen identifies CSDE1 as a key regulator of hepatic *LDLR* mRNA decay *in vivo*, making it a promising target for heart disease.

**Graphical Abstract:** 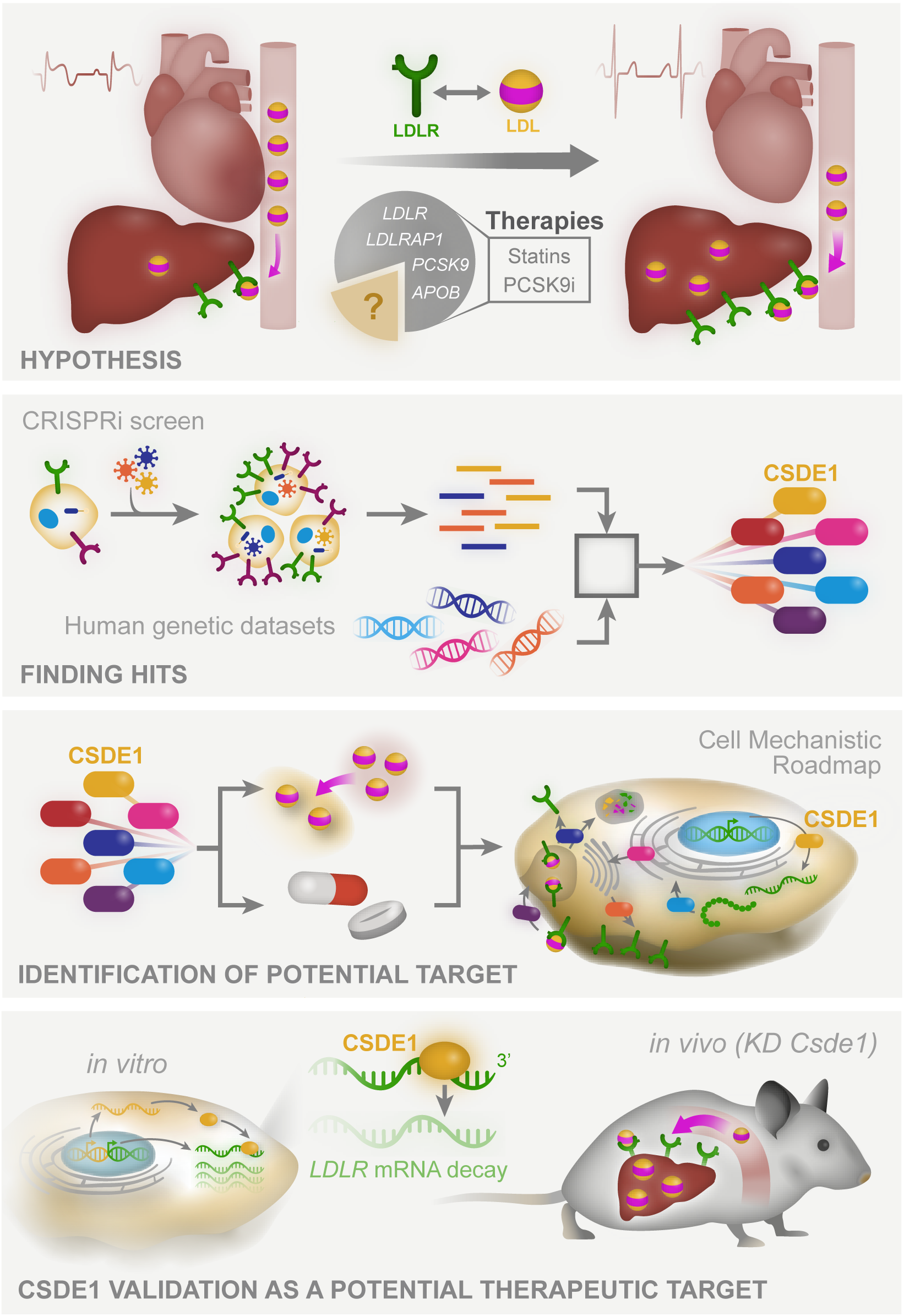

## Introduction

The low-density lipoprotein receptor (LDLR) delivers cholesterol from low-density lipoprotein (LDL) to cells to maintain membrane homeostasis (*1*). By clearing atherogenic LDL from the bloodstream, the hepatic LDLR protects against atherosclerotic heart disease (*2, 3*). Despite successful therapies that upregulate the hepatic LDLR and reduce heart attacks (*4*), cardiovascular disease remains the leading cause of death in Western countries (*5*). Lowering LDL beyond the levels achieved by HMG-CoA reductase inhibitors (statins) improves clinical outcomes without adverse effects (*6, 7*). Though there is a theoretical level at which LDL could get too low (*8*), this has yet to be discovered in large randomized trials (*9, 10*). Whether other LDLR regulatory mechanisms could be leveraged to further treat heart disease remains unknown.

The genetics of familial hypercholesterolemia (FH), which manifests as an isolated elevation in serum LDL, underlies the clinical success of LDLR upregulation by statins and PCSK9 inhibitors. Estimates suggest that 20-40% of FH phenotypes remain unexplained outside of the four major causes: *LDLR*, *APOB*, *PCSK9*, and *LDLRAP1* (*11, 12*). This implies that additional regulators of the LDLR exist. Advances in forward genetics (*13–15*) can now enable searches for tissue and disease-specific effects across the entire genome that may elude the sporadic natural variants found in observational studies, which themselves require compatibility throughout the entire lifespan and in all cell types. Moreover, hepatic delivery of gene silencing agents is effective in the clinic (*16*), providing a therapeutic modality against hits whose phenotypes are driven by expression in the liver. We therefore employed a genome-wide CRISPR interference screen for factors involved in hepatic LDLR regulation, both to understand the biology of this important receptor and to uncover potential therapeutic targets in cardiovascular disease.

## Results

### A Genome-Wide CRISPR Interference Screen for LDL Receptor Regulation

We engineered the HepG2 cell line, which models the regulation of the LDLR (*17*), to constitutively express a dCas9-KRAB fusion protein, enabling the knockdown of any given gene with an appropriate sgRNA (Fig. 1A) (*13, 14, 18*). Because statins (*19, 20*) and PCSK9 inhibitors (*21–23*) increase cell surface LDLR, we scored surface LDLR levels. To focus on factors that preferentially affect the LDLR over other receptors, we performed a parallel screen for regulators of the transferrin receptor (TFR). This critical player in iron metabolism shares a clathrin-mediated intake mechanism, but is otherwise orthogonally regulated from the LDLR (*24, 25*). Prior to our screen, we confirmed both dCas9-KRAB activity and an appropriate dynamic range for both LDLR and TFR regulation by transduction with sgRNAs expected to alter receptor levels in either direction (Fig. S1) (*26, 27*).

**Figure 1:**
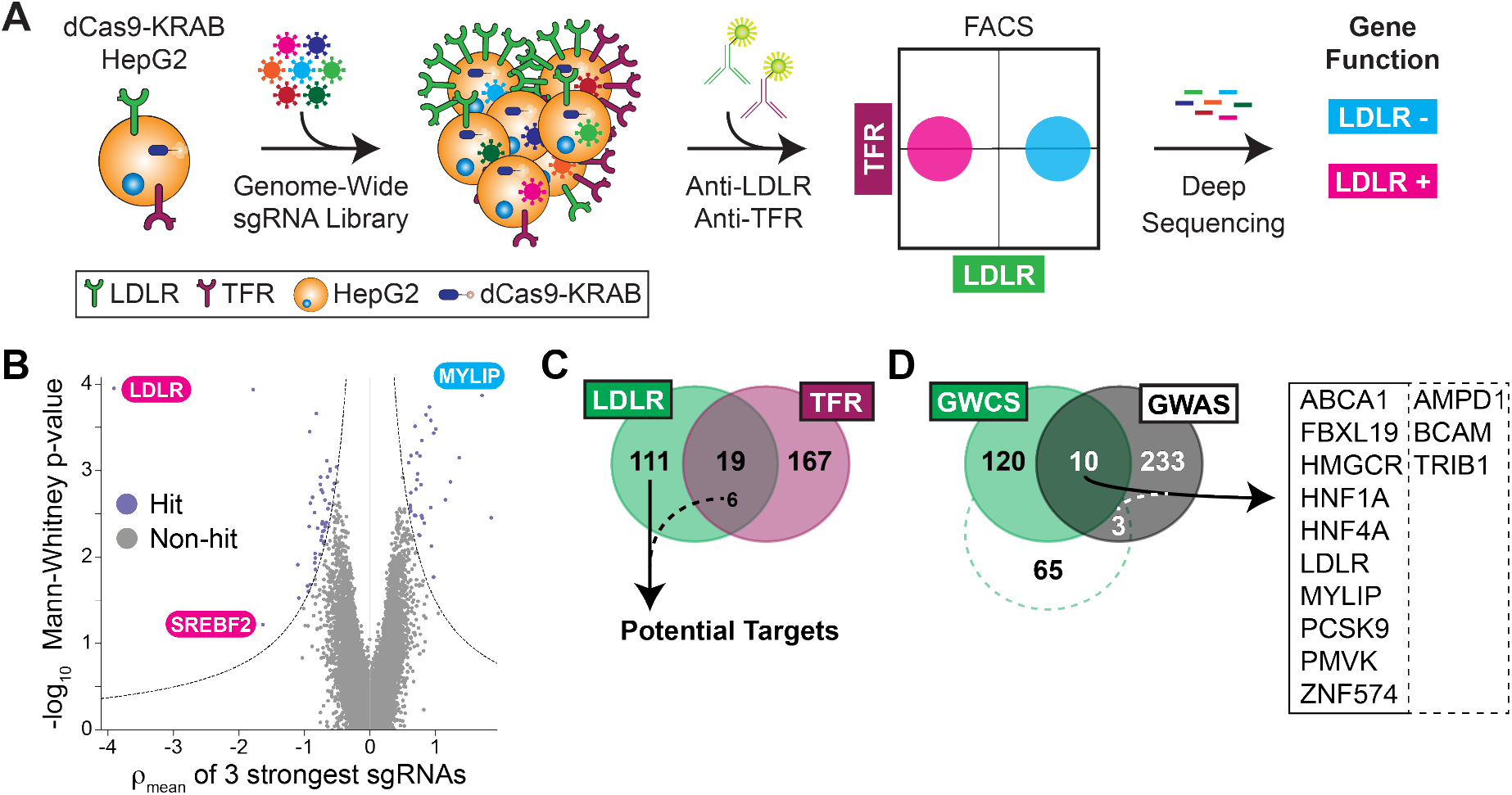
Genome-Wide CRISPR Interference Screen. *A)* Overall schematic of selection. See text for details. *B)* Volcano plot showing the statistical significance (Mann-Whitney test) of the guides recovered for each gene against the mean ρ phenotype of the three guides with the strongest effect. ρ is defined as the log_2_-fold enrichment for high LDLR expressing cells to the low LDLR expressing cells. Guides targeting known regulators of the LDLR are noted. *C)* Venn diagram showing the overlap between parallel LDLR and TFR screens. 6 guides common to both had opposing expression phenotypes in the respective screens and were included as specific hits. *D)* Venn diagram of hits between the LDLR screen (GWCS) and putative genes correlated with serum LDL from GWAS. The dotted line indicates a relaxed threshold for hit selection from LDLR screen, with only an additional 3 genes in the overlap. Overlap genes shown at right.

We next performed our pooled screens in parallel by transducing a library encoding sgRNAs with 5-fold coverage of the entire protein-coding human genome (*14*). We then selected the cells at the upper and lower third of receptor abundance by FACS and quantified the sgRNAs for each population via deep sequencing (Fig. S2, Tables S1-S4). We compared the degree of enrichment of LDLR or TFR surface levels in the high abundance to the low abundance cells (defined as ρ, Fig. 1B). We also compared the high and low receptor abundance cells to the unsorted population (defined as τ or γ, respectively) and included these results in our final hit count. This resulted in 130 total hits for the LDLR and 186 hits for the TFR (Tables S5-S6). We hypothesized that hits with shared phenotypes would likely have global effects on surface receptors, leaving us with 117 hits specific for LDLR regulation (Fig. 1C, Table 1). Gene ontology (GO) analysis (*28*) revealed a 15-fold enrichment for cholesterol metabolism as a biologic process (11 total hits, p = 5.7×10^-10^), providing confidence that we recapitulated our target biology. The hits also included 48 members of potentially druggable protein classes, including 29 with proposed enzymatic activity, and 22 hits were unclassified in GO databases (Fig. S3A).

**Table 1:**
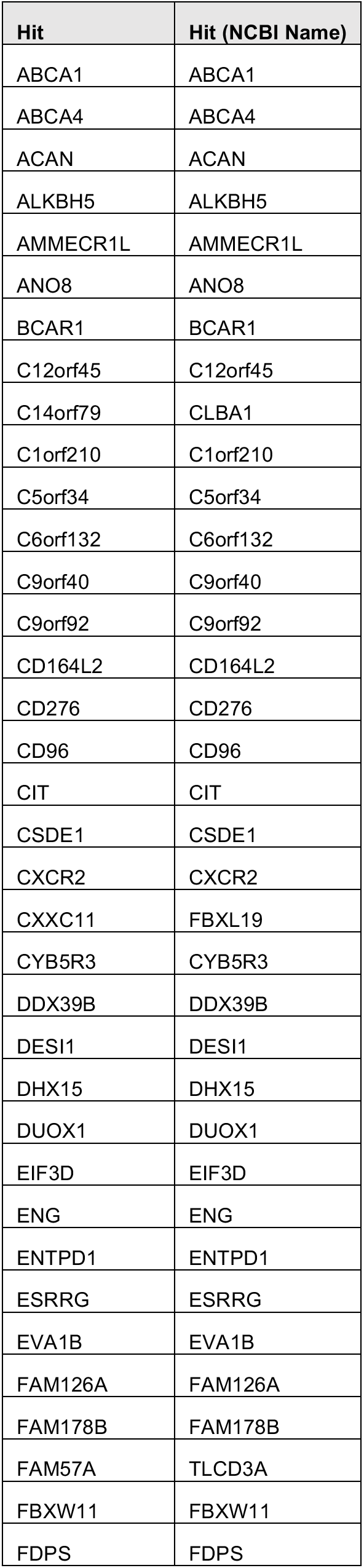
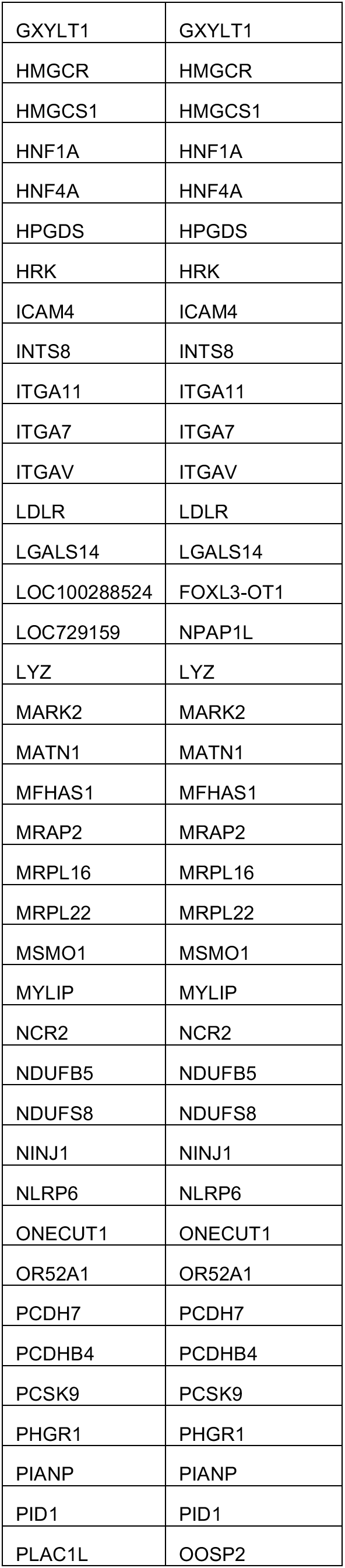
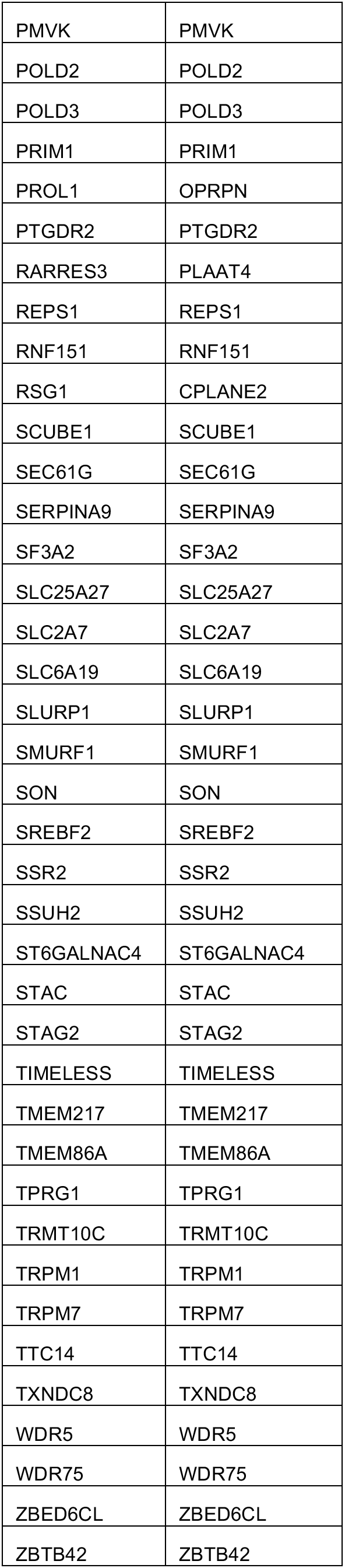
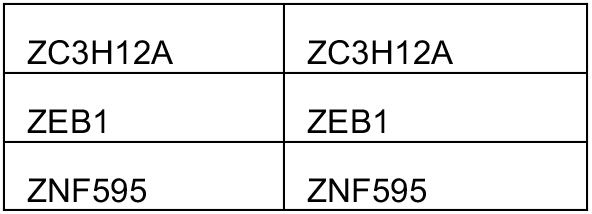
LDLR Specific CRISPRi Screen Hits. Hits are listed both by gene name in the genome-wide library(*14*) as well as NCBI name.

### Cross-referencing Human Genetic Datasets Identifies LDLR Regulators in vivo

We next compared genes associated with serum LDL cholesterol (LDL-C) from published genome-wide association studies (GWAS) (*29–31*) to our list of hits. Intriguingly, only 13 of these genes overlapped with our results (Fig. 1D), even when we relaxed our threshold for hit selection. To improve power for multiple hypothesis testing across the entire genome, we turned to 390,375 UK Biobank participants with genome-wide genotypes and known plasma lipids (Table S7) to search for variants associated with LDL-C amongst only our hits (*32*). We filtered to nonsynonymous protein coding variants in these hits by a threshold minor allele frequency (>0.001) and minimum statistical significance (Table 2). For *BCAM*, we found both an association between higher LDL-C and a nonsense variant, along with bidirectional associations between LDL-C and missense variants, suggesting that this pathway may be tunable. We also found associations between elevated LDL-C and variants in *MSMO1*, *C6orf132*, *HNF4A*, and *TIMELESS*, suggesting that these hits may be functional in the human and warrant further evaluation. The results also suggest that the accessible “genomic space” of the CRISPRi and GWAS strategies is only partially overlapping.

**Table 2:** Association of Nonsynonymous Variants in CRISPRi Screen Hits with Serum LDL-C in the UK Biobank. BETA indicates the linear regression standardized effect size, and P_BOLT_LMM indicates the linear mixed model p value using BOLT-LMM(*97*).

| GENE | Variant rsID | BETA | P_BOLT_LMM | Consequence | IMPACT |
| --- | --- | --- | --- | --- | --- |
| HNF4A | rs1800961 | 0.0564144 | 0 | missense_variant | MODERATE |
| BCAM | rs28399659 | -0.0174111 | 7.70E-29 | missense_variant | MODERATE |
| BCAM | rs200398713 | -0.0803165 | 1.80E-28 | splice_region_variant,intron_variant | LOW |
| BCAM | rs199922856 | -0.342179 | 6.20E-28 | missense_variant | MODERATE |
| BCAM | rs28399654 | 0.220592 | 6.10E-10 | missense_variant | MODERATE |
| BCAM | rs3810141 | 0.020077 | 5.50E-07 | stop_gained | HIGH |
| TIMELESS | rs2291738 | 0.00388393 | 0.00014 | splice_region_variant,intron_variant | LOW |
| BCAM | rs149302547 | -0.147327 | 0.005 | missense_variant | MODERATE |
| BCAM | rs1135062 | -0.0213642 | 0.0074 | missense_variant | MODERATE |
| C6orf132 | rs55772414 | 0.0116856 | 0.013 | missense_variant | MODERATE |
| MSMO1 | rs142496142 | 0.0432195 | 0.015 | missense_variant | MODERATE |

### Regulators of Surface LDL Receptor Abundance Affect Functional Uptake of LDL

To validate our screen results, we generated CRISPRi HepG2 cells harboring either of the two top-scoring sgRNAs for 77 of our hits as well as established controls. We preferentially tested hits with an increase in surface LDLR upon inhibition, as well as those with potentially druggable functions or lacking associated GO terms. Since surface receptor levels might not necessarily correlate to increased function, we evaluated both LDLR and TFR surface phenotypes alongside a functional assay of LDL uptake (*33*). Lastly, as knockdowns could also cause growth phenotypes, we assayed the number of cells surviving to FACS analysis as a proxy for viability.

We recapitulated the phenotypes for receptor abundance for at least one of the guides in the majority of the hits (55 genes, 71% of those tested, Table S8). Moreover, for 40 of these genes, both sgRNAs independently validated, suggesting against an off-target effect. We visualized these hits based on their effects, at single cell resolution, on LDLR and TFR levels, the LDLR/TFR ratio, functional LDL uptake and number of cells surviving to analysis (Figs. 2, S3B). Notably, most knockdowns had independently validated effects on LDLR abundance and LDL uptake of similar or greater magnitude than our *HMGCR* or *PCSK9* controls.

**Figure 2:**
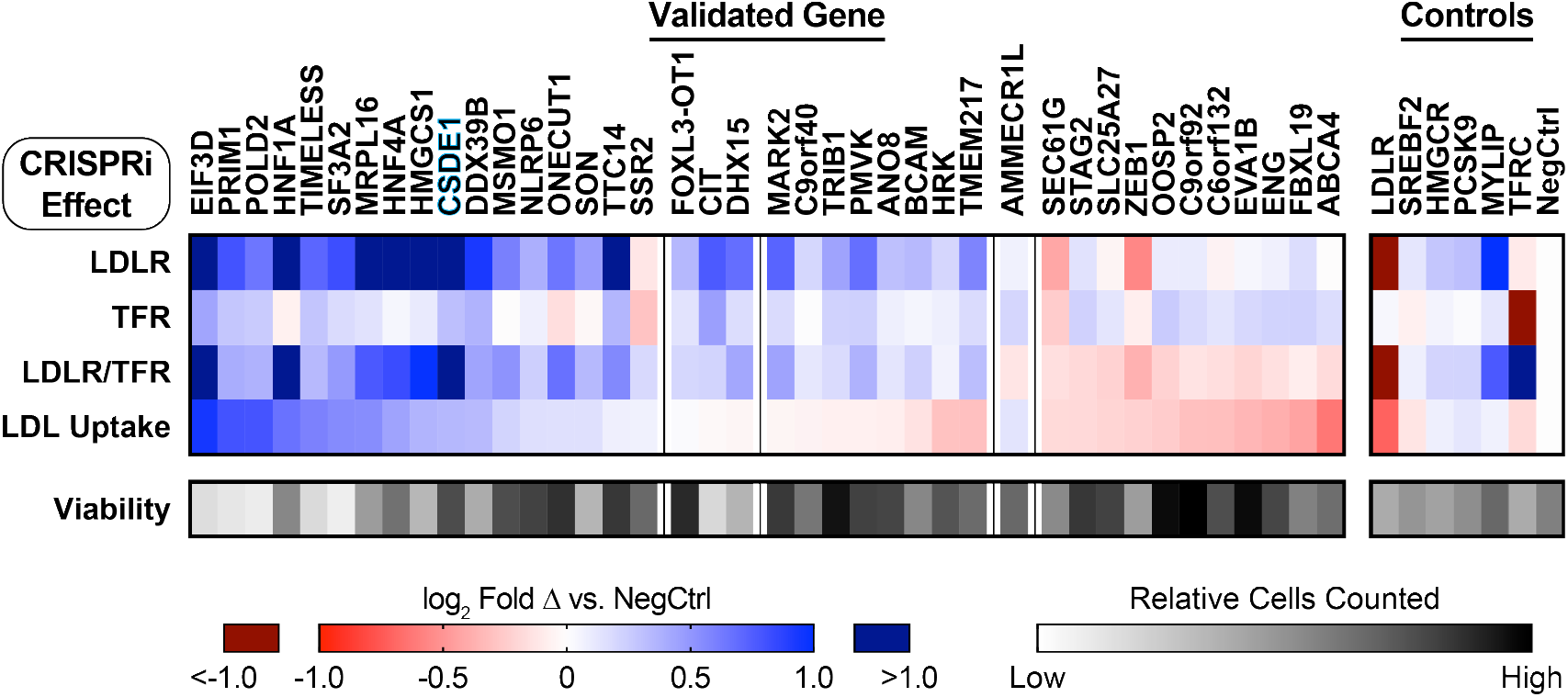
Validation of LDLR CRISPRi Hits. Heatmap showing receptor abundance (LDLR, TFR, and LDLR/TFR ratio) and function (LDL uptake) for dCas9-KRAB HepG2 cells transduced with sgRNA targeting the indicated gene, analyzed by flow cytometry. Hits are grouped according to directional effect on LDLR abundance, and then within groups, by effect on LDL uptake (with uptake from *FOXL3-OT1*, *CIT*, and *DHX15* sgRNAs not significantly different from negative control sgRNA). *CSDE1* highlighted in blue. Control sgRNAs shown at right. Readouts show log_2_ fold change compared to transduction with negative control sgRNA and represent the weighted average of the effects from both sgRNAs targeting each gene. Viability indicates the relative number of cells surviving to flow cytometry in the experiments. Functional classification of genes is shown visually in Supp. Fig. 3. Note that LDLR/TFR is a separately ascertained value from individual cells, and not a derived parameter from aggregate data. Only the hits for which two separate sgRNAs independently validated for receptor expression are shown, defined as p < 0.05 via Holm-Sidak corrected T-test. Data represent summary information from 3 to 4 independent experiments.

Knockdown of hits expected to alter cellular cholesterol balance or transcriptionally regulate the LDLR showed directionally consistent effects between LDLR abundance and function (Fig. 2). For genes in the enzymatic pathway of cholesterol metabolism (*34*) (*HMGCS1* and *MSMO1*), this is consistent with activation of SREBP2-mediated *LDLR* transcription. For genes encoding certain transcription factors (*HNF1A* (*35*), *HNF4A* (*36*), *ONECUT1* (*37*), and *ZEB1* (*38*)), this is consistent with an effect on *LDLR* transcription itself. Knockdowns of *SLC25A27*, which encodes a mitochondrial uncoupling protein (*39*), and *ABCA4*, encoding a known lipid transporter (*40*), both exhibited reductions in LDLR abundance and function (Fig. 2). These genes could plausibly induce a negative lipid balance, increasing LDL uptake via both LDLR dependent and independent mechanisms.

Targeting of hits that either affect multiple transcriptional pathways or regulate endocytosis showed opposite effects on LDLR abundance and function. Knockdown of *TRIB1*, a GWAS hit (*29*) encoding a pseudokinase that regulates the COP1 E3 ligase (*41, 42*) and affects multiple transcription factors (*43*), showed this phenotype. In the mouse, *TRIB1* overexpression lowers serum cholesterol, while the knockout has the opposite effect (*44, 45*), consistent with our results. Knockdown of *AP2M1*, a TFR screen hit that encodes an adaptor protein required for endocytosis (*46*), was similar, consistent with an accumulation of non-functional receptors at the cell surface. This phenotype, though specific to the LDLR, was also seen with knockdown of *BCAM*, which encodes a membrane cell adhesion molecule (*47*) identified by GWAS (*31*), and *TMEM217*, which encodes an uncharacterized transmembrane protein (Figs. 2, S4). This suggests that these proteins could have a similar endocytosis adaptor function specific for the LDLR, akin to *LDLRAP1* (*48*), in which mutations cause a recessive form of FH.

### Pharmacologic Inhibition of Clinically Relevant Pathways Provides Mechanistic Insight into Putative LDLR Regulators

We next turned to pharmacologic approaches to perturb specific pathways of LDLR regulation. We hypothesized that hits might alter cholesterol metabolism, LDLR recycling, or a yet unspecified pathway. By combining CRISPRi knockdown with either a statin, to inhibit endogenous cholesterol biosynthesis (*49, 50*), or a PCSK9 inhibitor, to arrest LDLR lysosomal degradation (*23*), and assessing the combined effect, we inferred mechanistic information about the target gene. Furthermore, we hypothesized that either additive or potentiating effects between a clinically validated therapy and a hit gene might suggest promising therapeutic targets.

We evaluated the receptor abundance and function phenotypes for 29 of our validated hits in the presence or absence of a statin (*51*) or PF-846, a selective inhibitor of PCSK9 translation (*52, 53*) (Fig. 3, Table S9). We calculated a synergy score by subtracting the differential effects of CRISPRi knockdown, compared to the control, in the presence of compound from that with the DMSO vehicle. A more positive value indicated synergy, and a more negative value indicated antagonism.

**Figure 3:**
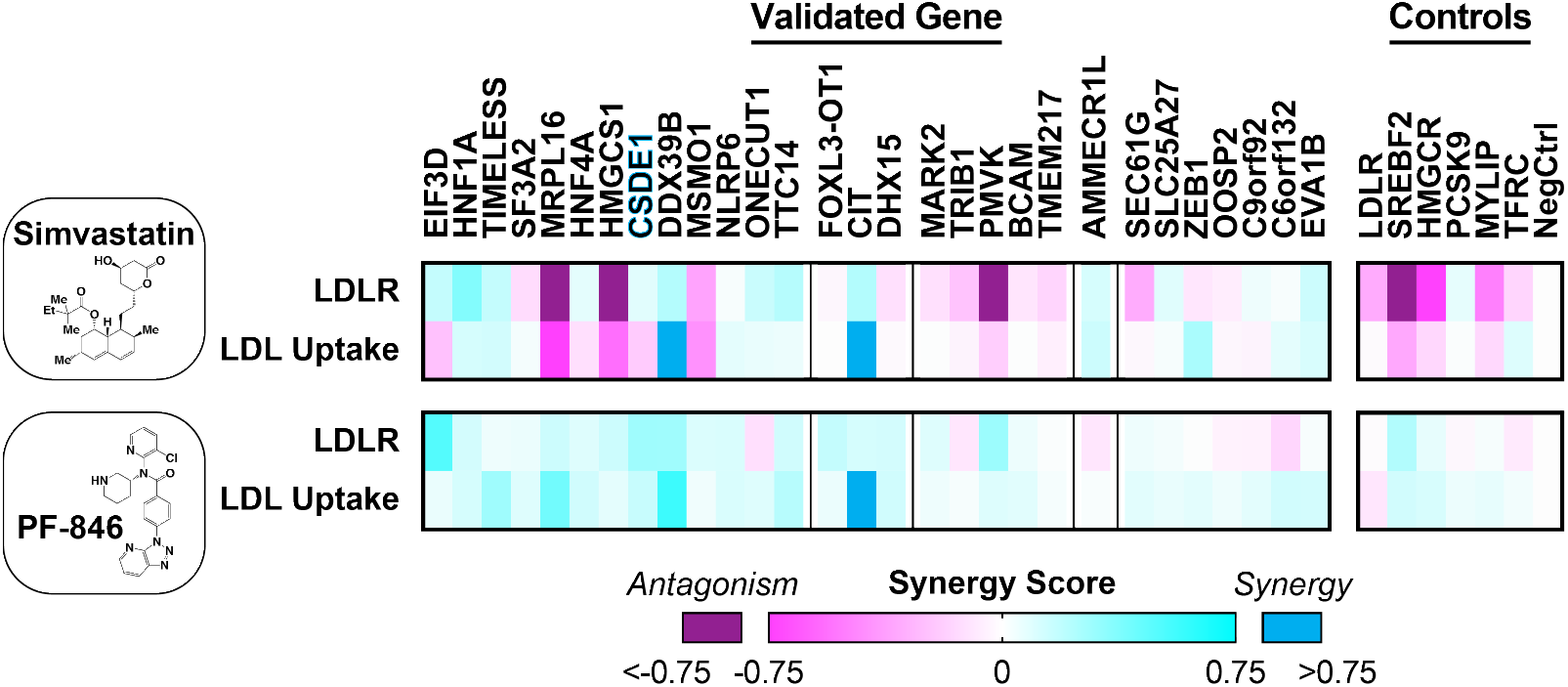
CRISPRi Knockdown Synergy with Statin and PF-846. Heatmap showing synergy score with statin (top) or PF-846 (bottom) for knockdowns of indicated validated genes with a single sgRNA for separate LDLR abundance and function experiments. Hits are grouped first according to overall effect on LDLR abundance, and secondarily by effect on LDL uptake, as in Fig. 2. *CSDE1* highlighted in blue. Data represent summary information from 4 independent experiments.

Upon knockdown, regulators of cholesterol biosynthesis (*SREBF2*, *HMGCR*, *HMGCS1*, *MSMO1*, and *PMVK*) showed antagonism with the statin, but mild synergy with PCSK9 inhibition (Fig. 3). This is consistent with the SREBP2-mediated *PCSK9* transcription that underlies the clinical synergy between statins and PCSK9 inhibitors. The synergy phenotypes for knockdown of *MRPL16*, which encodes a structural component of the mitochondrial ribosome (*54*), mirrored these cholesterol biosynthetic genes (Fig. 3), suggesting that MRP-L16 may play a role in the mitochondrial generation of metabolic precursors to sterol biogenesis. In contrast, *C6orf132* knockdown showed the opposite phenotype: mild synergy with a statin, and mild antagonism with PF-846 (Fig. 3). Given that C6orf132 localizes to the Golgi (*55*), this suggests it may function by facilitating LDLR delivery to the cell surface, prior to any interaction with extracellular PCSK9. For some transcription factors, the synergy phenotypes can point to their downstream targets. For example, synergy of *HNF1A* knockdown with a statin (Fig. 3) is consistent with disruption of HNF1-α-mediated *PCSK9* transcription (*56*).

### CSDE1 Regulates the Stability of LDLR mRNA

One of our strongest hits, *CSDE1*, also known as upstream of N-ras (UNR), encodes an RNA binding protein with varied regulatory functions (*57–59*), including mRNA decay (*60*). As the *LDLR* 3’ UTR consists of adenylate-uridylate (AU)-rich elements (AREs) implicated in mRNA stability (*61*), we hypothesized that CSDE1 could mediate the degradation of the *LDLR* transcript, thereby explaining its observed receptor abundance, function, and synergy phenotypes.

In the setting of *CSDE1* knockdown, we observed progressively higher LDLR abundance with sterol depletion and a concomitant statin (Figs. 4A, S5A-C). This suggests the mechanism of *CSDE1* disruption is at least additive with SREBP2-mediated *LDLR* transcription and statin therapy. We also observed that overexpression of isoform 1 of CSDE1, but not isoforms 2 through 4, reduced surface LDLR in HepG2 cells (Figs. 4B, S6A-D). Overexpression of all four isoforms of CSDE1 downregulated LDLR levels in the *CSDE1* knockdown cells, though isoform 1 showed the strongest effect (Fig. S6E-I). The opposing directional effects of *CSDE1* knockdown and overexpression suggest that, under physiologic expression conditions, isoform 1 of CSDE1 is a rate-limiting regulator of the LDLR.

**Figure 4:**
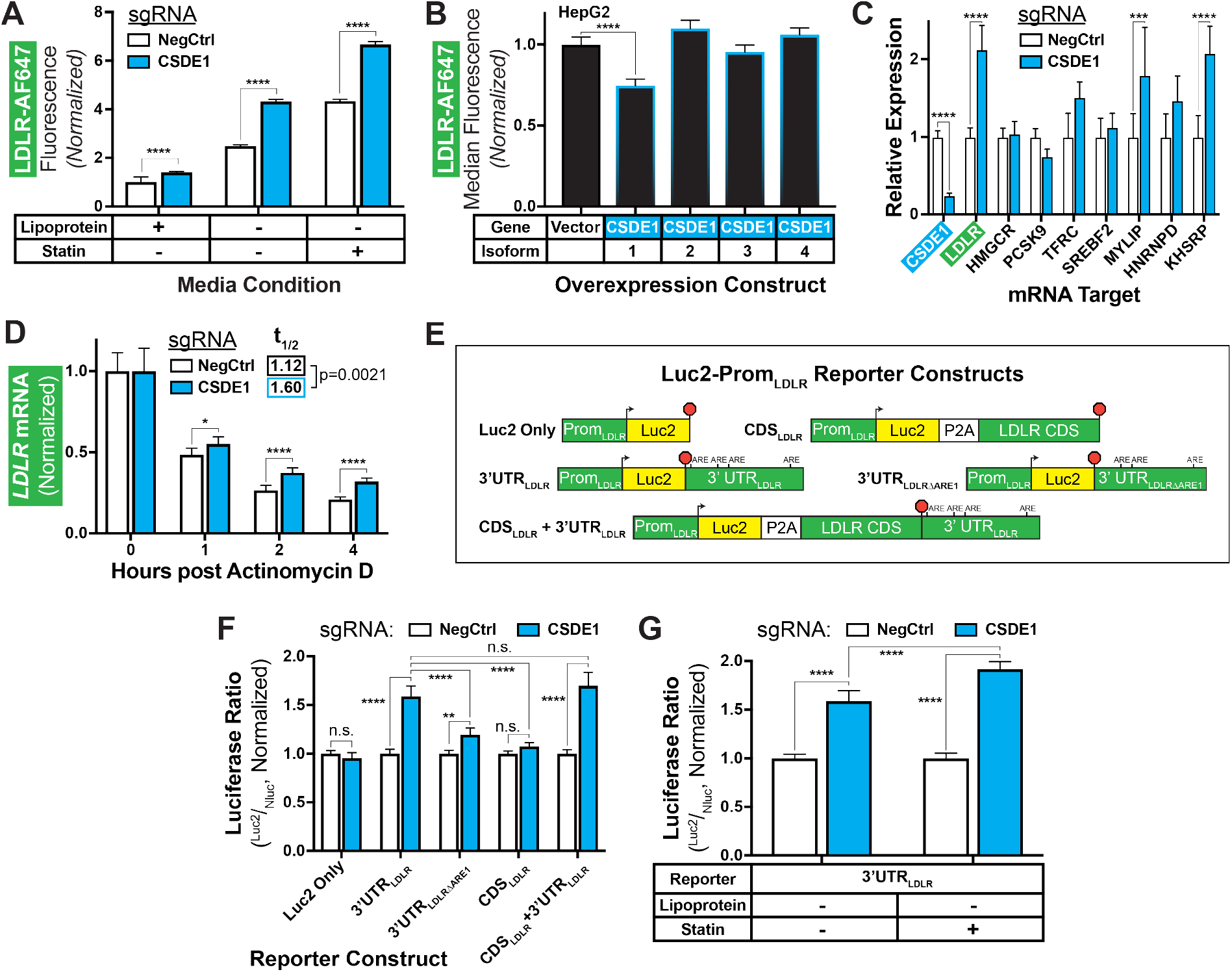
CSDE1 Mediates *LDLR* mRNA Decay. *A)* Relative LDLR abundance, by normalized mean fluorescence, in engineered dCas9-KRAB HepG2 cells transduced with indicated sgRNAs (CRISPRi cells) and grown in the indicated media conditions. *B)* Relative LDLR abundance, by normalized median fluorescence, in HepG2 cells overexpressing indicated CSDE1 isoforms. One-way ANOVA with Tukey’s multiple comparisons test shown. *C)* Relative expression, by qPCR, of indicated mRNA targets in CRISPRi cells under sterol-depleted conditions. *D)* Relative expression, by qPCR, of *LDLR* mRNA in CRISPRi cells after arrest of transcription with actinomycin D. Data are normalized to results at T=0 within the sgRNA evaluated to illustrate the change in time. t_1/2_ shown indicates the best fit data to a one-stage exponential decay equation. Unpaired t-tests with Holm-Sidak correction shown for pairwise comparisons. Extra sum-of-squares F test shown for decay equation. *E)* Schematics of Luc2-Prom_LDLR_ reporter constructs, illustrating *LDLR* promoter, start site (arrowhead), P2A ribosomal skipping sequence (if present), AREs in 3’ UTR, stop codon (red octagon), and indicated regions of the *LDLR* gene. ARE = Adenylate-uridylate (AU) rich element. Schematics not to scale. *F)* Ratiometric luciferase outputs, normalized to negative control, of CRISPRi cells transfected with indicated luciferase constructs. *G)* Ratiometric luciferase outputs of 3’ UTR addended luciferase constructs in CRISPRi cells. *All panels)* Error bars indicate 95% confidence intervals. Data represent summary information from 3-4 independent experiments. n.s. = not significant, * = p < 0.05, ** = p < 0.01, *** = p < 0.001, and **** = p < 0.0001. Unless otherwise indicated, two-way ANOVA with Sidak’s multiple comparisons test was used for statistical analysis.

Consistent with our mechanistic hypothesis, we noted over a 2-fold increase in steady-state mRNA levels of *LDLR* (Fig. 4C), as well as depleted CSDE1 (Figs. 4C, S7), in the *CSDE1* knockdown cells. Among control mRNA targets, we also observed significant increases in *MYLIP* and *KHSRP* mRNA. These gene products downregulate the LDLR (*26, 62*), which is the opposite of our observed phenotype, suggesting that the direct effect of *CSDE1* knockdown on the *LDLR* mRNA predominates. To specifically evaluate transcriptional decay, we treated cells with actinomycin D and measured *LDLR* transcript levels over time. We observed significantly higher *LDLR* mRNA in the *CSDE1* knockdown cells at all subsequent timepoints (Fig. 4D). The mRNA half-life, modeled by a single-phase decay equation, was nearly 1.5-fold longer for the *CSDE1* knockdowns compared to controls (p = 0.0021, Fig. 4D). Notably, *CSDE1* knockdown had no significant effect on *HMGCR*, *SREBF2*, *PCSK9*, or *TFRC* mRNA levels over time (Fig. S8), suggesting that the effect on mRNA stability was isolated to *LDLR* among our tested transcripts.

To probe the relationship of CSDE1 to the *LDLR* 3’ UTR, we transiently expressed luciferase constructs (Fig. 4E) under control of the native *LDLR* promoter in the *CSDE1* knockdown cells. The luciferase-only constructs showed appropriate physiologic upregulation by sterols, regardless of *CSDE1* knockdown (Fig. S9). Constructs fused to the *LDLR* 3’ UTR, but not those fused to the *LDLR* coding sequence alone, exhibited increased reporter activity with *CSDE1* knockdown (Fig. 4F). Notably, this increase in activity was attenuated by removing the first of four AREs (*61, 63*) from the 3’ UTR (Fig. 4F). Activity of the 3’ UTR-fused construct increased further with statin coadministration (Fig. 4G), suggesting that *CSDE1* knockdown may be synergistic with statins, consistent with our prior results (Fig. 3). Taken together, we conclude that under physiologic conditions, CSDE1 mediates decay of the *LDLR* mRNA through its 3’ UTR, with the first ARE of the UTR required for its full effect.

### Disruption of CSDE1 Upregulates the LDLR in vivo

We then turned to an *in vivo* model in zebrafish, as the 3’ UTR of its ortholog *ldlra* (XM_005163870.4) is highly AU-rich and contains at least two canonical ARE sequences for mRNA regulation (*64*). We employed yolk microinjection of a Cas9-ribonucleoprotein (RNP) complex containing redundant guides to achieve near-saturation gene disruption (*65*), followed with dietary cholesterol supplementation, and evaluated total cholesterol in the larvae (*66*). Targeting of *csde1* protected against cholesterol accumulation, with a modest (12%) but significant reduction in total cholesterol in 8-day post fertilization (dpf) zebrafish, without any obvious phenotypic abnormalities (Figs. 5A, S10). By contrast, targeting of *ldlra* showed the expected 1.4-fold increase in total cholesterol (Fig. 5A) (*66*).

**Figure 5:**
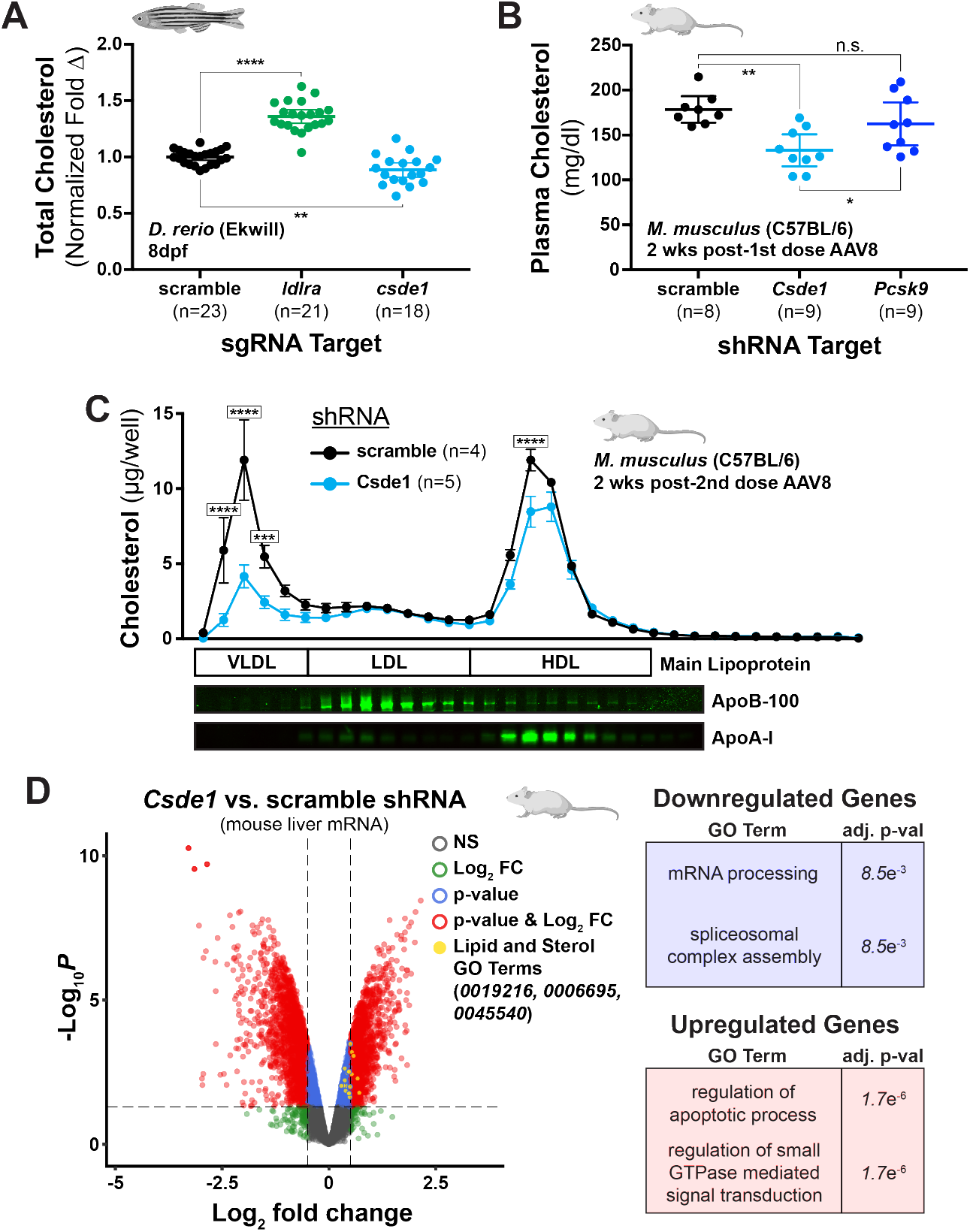
CSDE1 Disruption Upregulates the LDLR *in vivo*. *A)* Total cholesterol, in µg cholesterol per mg of total protein, of homogenates of 8 days post fertilization (dpf) zebrafish larvae fed a high-cholesterol diet and subjected to Cas9 mediated gene disruption of indicated target. Data are normalized to the scramble control of a particular experiment. Each point represents a homogenate consisting of 10 larvae. Data represent summary information from 4 independent experiments. One-way ANOVA with Holm-Sidak’s multiple comparisons test shown. *B)* Total fasting plasma cholesterol of C57BL/6 mice on an atherogenic diet, 2 weeks after transduction with AAV8-packaged shRNA against indicated target. Each point represents an individual mouse. One-way ANOVA with Tukey’s multiple comparisons test shown. Error bars = 95% confidence intervals. *C)* Cholesterol levels of fractions collected from gel filtration of plasma from individual mice, harvested 2 weeks after transduction with second dose of AAV8-packaged shRNA. Note that fractions shown begin with the elution front from the size-exclusion column. Each dot represents the mean cholesterol level from mice in the same intervention arm. Immunoblots from representative fractions against mouse ApoB-100 and ApoA-I shown below. Two-way ANOVA with Sidak’s multiple comparisons test shown to illustrate comparison between treatment arm within a given fraction. Error bars = standard error of the mean. *D)* Volcano plot showing differentially expressed genes between *Csde1* and scramble shRNA treatment arms, filtered for effects of viral transduction. Statistical significance is shown on the y axis and strength of effect is shown on the x axis. Comparison made among the 3 mice in each arm with the highest *eGFP* transcript expression, as a proxy for transduction efficiency. Genes reaching threshold significance for p-value (blue), log_2_ fold change (green), both (red), or neither (grey) annotated accordingly. Genes associated with lipid and sterol metabolic GO terms highlighted in yellow (0019216 = lipid metabolic process, 000695 = cholesterol biosynthetic process, and 0045540 = regulation of cholesterol biosynthetic process). Leading upregulated and downregulated GO terms of all statistically significant differentially expressed genes noted in the boxes at right. *All panels)* n.s. = non-significant, * = p < 0.05, ** = p < 0.01, *** = p < 0.001, and **** = p < 0.0001.

We next probed the effect of *Csde1* gene silencing in the mouse as a therapeutic proof-of-principle, given even greater homology between the 3’ UTRs of the murine and human *LDLR* orthologs (*67*). Using C57BL/6 mice on an atherogenic diet (*68*), we delivered shRNA against *Csde1* (*59*), *Pcsk9*, or scramble control via low-dose adeno-associated virus 8 (AAV8). Two weeks later, we observed a 25% reduction in fasting plasma cholesterol in the *Csde1* knockdown mice, which exceeded the effect of *Pcsk9* knockdown (Fig. 5B). We then re-dosed the *Csde1* and scramble AAV8-shRNA and, 2 weeks later, observed an even stronger phenotype (Fig. S11A). Lipoprotein fractionation of the mouse plasma showed that *Csde1* knockdown mostly affected the VLDL-containing fractions (Fig. 5C), consistent with upregulation of the murine LDLR on our dietary background (*69, 70*). Accordingly, we observed an increase in *Ldlr* expression in the liver tissue of the *Csde1* knockdown mice (Fig. S11B). Notably, we observed no differences in appearance or behavior of the mice, nor in plasma levels of alanine aminotransferase activity (Fig. S11C), arguing against hepatic or systemic toxicity of either the *Csde1* shRNA or the gene knockdown.

To gain further insight, we performed bulk RNA sequencing on the liver tissue (Figs. 5D, S11D-F, Table S10). We compared the *Csde1* knockdown to control (scramble) mice, using the mice with the highest transcript counts of a vector-delivered eGFP reporter to control for variations in transduction efficiency. We then filtered our results for the differentially expressed transcripts in the control mice at the extremes of eGFP expression, to control for the effects of viral transduction alone. As expected, we found higher *Ldlr* expression in the *Csde1* knockdown mice (log_2_FC = 0.43, p = 0.0029, Table S10). Consistent with our mechanistic hypothesis, GO enrichment analysis of the differentially expressed genes revealed that mRNA processing was the most significantly downregulated biological process in the *Csde1* knockdown mice (OR = 2.7, adj. p = 0.0085, Figs. 5D, S11F, Table S11). Spliceosome complex assembly was also significantly downregulated (OR = 7.4, adj. p = 0.0085, Figs. 5D, S11F, Table S11).

## Discussion

The powerful biology of the LDLR is unquestioned in cardiovascular medicine (*71*). Since their introduction, statins, which upregulate the LDLR, have become a major public health success, and with the discovery of PCSK9 and the therapeutic antibodies targeting it, patients can safely reach much lower LDL levels than is achievable by statins alone (*10*). Together, this suggests that we can push further on this LDL-LDLR axis and still achieve a clinical benefit.

In this study, we modeled a clinically relevant phenotype of LDLR abundance and function, complementing the independent investigations of other groups (*72, 73*). When synthesizing our screening and validation data together with large-scale genomics and additional pharmacologic perturbations, we produce an exploratory map of potential regulatory mechanisms for the LDLR (Fig. 6). These data represent not just promising targets but also pathways likely to be impacted by therapies already in use in the clinic.

**Figure 6:**
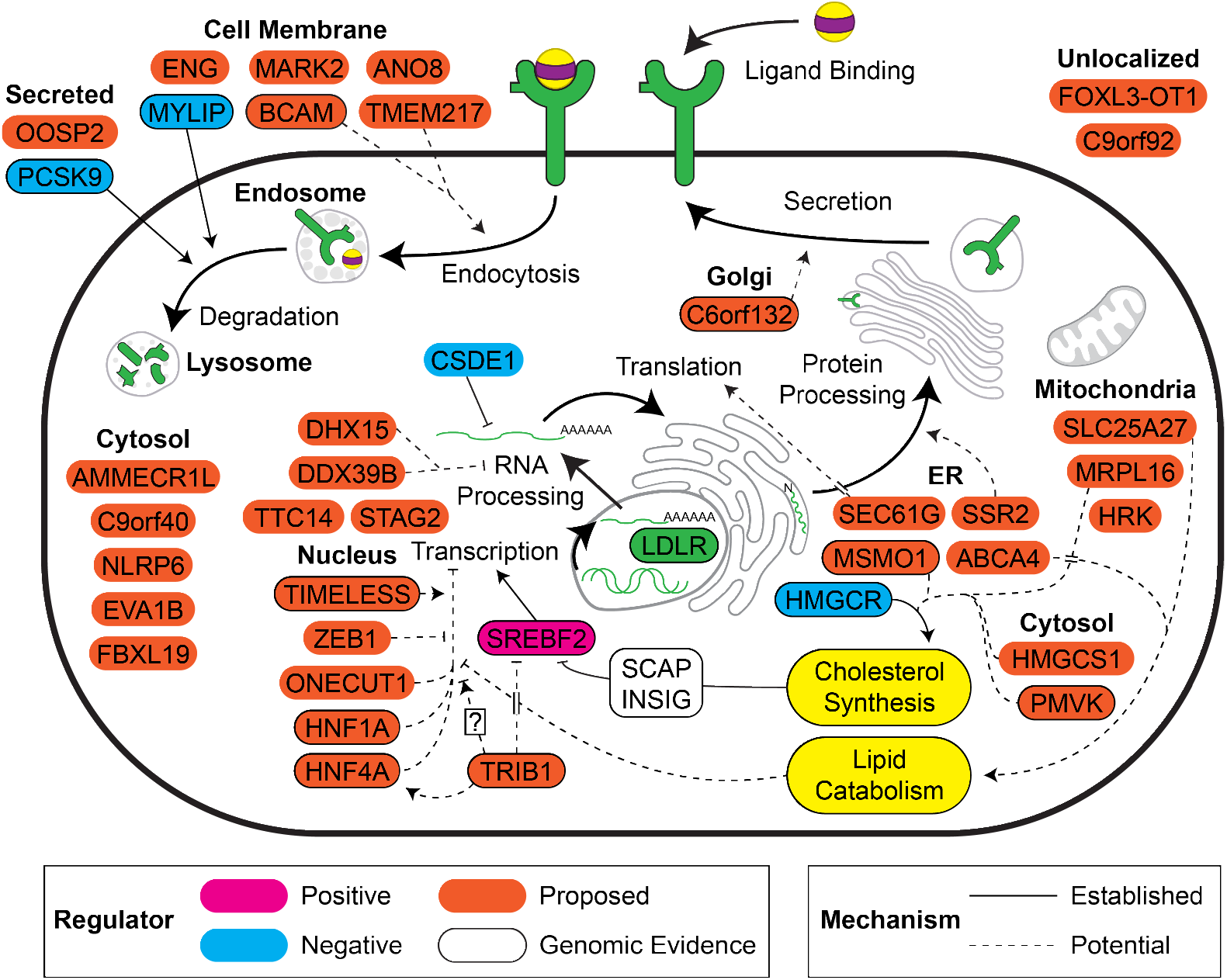
An Exploratory Map of Potential LDLR Regulatory Targets. Genes identified and validated in the screen are mapped by cellular localization and possible mechanisms of effect. Known downregulators are shown in cyan (including CSDE1 given the results presented in the current study) and known upregulators shown in magenta. Validated hits with observed effects on cell proliferation or viability are excluded.

We have shown that *CSDE1*, one of our strongest hits, regulates LDLR levels in HepG2 cells by promoting *LDLR* mRNA decay via its 3’ UTR. These data lay in concert with CSDE1’s destabilizing effects on other transcripts, such as c-Fos (*60*). We have also shown that *in vivo* knockdown of *Csde1* upregulates the hepatic LDLR and improves atherogenic lipid profiles in mice. This mimics the effect of deleting the 3’ UTR *in vivo* (*74*) and illustrates the promise of targeting CSDE1 to lower LDL and protect against atherosclerosis. It is notable that several small molecules, including triciribine (*63*) and berberine (*67, 75*), have stabilizing effects on *LDLR* mRNA, though whether their mechanisms directly involve CSDE1 remain to be elucidated. The magnitude of LDLR upregulation imparted by *CSDE1* knockdown mirrors or exceeds that of *HMGCR* and *PCSK9* in both tissue culture and mouse models, suggesting that a high-fidelity approach targeting CSDE1-mediated *LDLR* mRNA decay in the clinic could have similarly impressive effects. Additionally, our mechanistic data suggest that targeting CSDE1 would be at least additive with the use of statins.

The degree to which CSDE1 inhibition affects other transcripts, or other tissues (*59, 76*), remains an important question. As an RNA chaperone, CSDE1 can have a variety of effects, from mRNA stabilization (*58*) to promotion or inhibition of translation (*77–80*), dependent on the identity of the RNA it binds and the cofactors with which it interacts. Intriguingly, though CSDE1 was found to bind biotinylated *LDLR* 3’ UTR transcripts in HepG2 cell lysates (*62*), cross-linking immunoprecipitation approaches in both mouse brain and melanoma cells failed to identify *LDLR* mRNA as a CSDE1 binding partner (*81, 82*). This suggests that the CSDE1-*LDLR* interaction is context dependent. Advances in liver-specific delivery of gene-silencing agents (*16, 83*), novel gene editing technologies (*84*), and small molecules (*85*) offer the possibility that selectively targeting hepatic CSDE1 for cholesterol lowering could avoid systemic toxicities.

Though we observed no toxicity, transcriptional profiling suggests that disrupting hepatic CSDE1 upregulates both apoptosis-related and GTPase-mediated signaling pathways (Figs. 5D, S11F, Table S11). Notably, CSDE1 appears to protect from apoptosis in both colorectal cancer (*86*) and Huh7 cells (*87*). Though biologically plausible, we hesitate to make definitive conclusions from these data since the cholate-rich diet we used to obtain hyperlipidemia also causes liver inflammation and hepatic steatosis (*88*). Despite this confounder, we expect that our transcriptomic analysis will guide further investigations of potential toxicities from hepatic CSDE1 disruption. Given that CSDE1 has such varied effects on other transcripts, future mechanistic dissection of the hepatic CSDE1-*LDLR* interaction could identify what makes this relationship unique and guide a potential therapeutic strategy. Combination therapies targeting interconnected pathways to disease can provide increased benefits without inducing extreme side effects, with angiotensin receptor blockade and neprilysin inhibition in heart failure a prominent clinical example (*89*). Though speculative, we are intrigued by the effects of CSDE1 on mRNA splicing (Figs. 5D, S11F, Table S11), and we note in particular that the spliceosome helicase DDX39B was both validated as a regulator of the LDLR from our CRISPRi screen (Fig. 2, Table S8) and downregulated by *Csde1* knockdown *in vivo* (log_2_FC = −0.32, p = 0.011, Table S10). To the extent hepatic CSDE1 utilizes specific factors to downregulate *LDLR* mRNA, simultaneous tissue-specific drugging of both CSDE1 and these factors could widen the overall therapeutic window. We anticipate that our study will serve as the seed for these and other further investigations.

## Methods

### Study Design

We designed the study as a discovery biology experiment to identify new regulators of the LDL receptor. We used an established tissue culture model, HepG2 cells, to evaluate for LDL receptor regulation. We used wild-type zebrafish (Ekwill) and wild-type mice (C57BL/6) to validate the contribution of our top hit, CSDE1, to LDL receptor regulation *in vivo*. We evaluated sufficient cells for the LDLR and TFR screens to provide adequate coverage for transduction and downstream sequencing of each sgRNA in the genome-wide library. Sample sizes for animal experiments were estimated to provide 80% power (two-tailed α = 0.05) for a 25% effect in cholesterol levels compared to controls, based on effects in these models in the existing literature. The numbers of animals used in each experiment are noted in the figures and manuscript. Unless otherwise noted, all *in vitro* data are representative of multiple (≥ 3) experimental outcomes to ensure robust outcomes. Experiments were not performed in a blinded fashion. All animal studies were performed in accordance with IACUC approved protocols at the University of California, San Francisco.

### Plasmids and Cloning

SFFV-dCas9-BFP-KRAB (Addgene 46911), CRISPRi/a v2 (Addgene 84832), pMD2.G, dR8.91, and the hCRISPRi v2 top5 sgRNA library (Addgene 83969) were gifts from L. Gilbert and J. Weissman. Oligonucleotides of the protospacers of validated sgRNA sequences (*14*), as well as those for PCR amplification and isothermal assembly, were obtained from Elim Biopharmaceuticals (Hayward CA). Protospacers were cloned into the CRISPRi/a v2 vector using restriction enzyme digest (BlpI and BstXI, ThermoFisher, Waltham MA) and ligation with 10× T4 ligase (NEB, Ipswich MA). CSDE1 overexpression constructs were created by PCR expansion of *CSDE1* (HsCD00949797, DNASU, Tempe AZ) or AcGFP1 (vector control, pIRES2-AcGFP1, Clontech, Mountain View CA) and isothermal assembly (*90*) into the pcDNA5/FRT/TO backbone (ThermoFisher), followed by site-directed mutagenesis to generate the four CSDE1 isoforms (UniprotKB O75534 1 through 4). pLuc2-Prom_LDLR_ was created by PCR expansion of the target luciferase from pGL4Luc-RLuc (Addgene 64034), custom gene synthesis of the LDLR promoter (NCBI Reference Sequence NG_009060.1, from −687 bp to the *LDLR* start codon, Twist Biosciences, South San Francisco CA) and isothermal assembly into the pcDNA5/FRT/TO backbone. pSS-NLuc was created by PCR expansion of the target luciferase from pNL1.1 (Promega, Madison WI) into a vector containing the PCSK9 signal sequence from the same backbone (*91*). The remaining pLuc2-Prom_LDLR_ constructs were created by PCR expansion of the coding region of *LDLR* (HsCD00004643, DNASU), custom gene synthesis of the 3’ UTR of the *LDLR* mRNA (NCBI Reference Sequence NM_001252658.1, Twist Biosciences), or custom oligonucleotides to add the P2A ribosomal skipping linker (*92, 93*) and isothermal assembly into pLuc2-Prom_LDLR_, as appropriate for each construct. All plasmids were confirmed by Sanger sequencing. Expansion of the top5 sgRNA library was as previously described (*13*).

### Cell Culture and Lentiviral Production

HepG2 (ATCC HB-8065) and derivatives were cultured in low-glucose DMEM (1 g/L, ThermoFisher) with 10% FBS (Axenia BioLogix, Dixon CA), GlutaMax (ThermoFisher) and 1× penicillin-streptomycin (ThermoFisher), and sent thrice through a 21g needle during passaging to minimize cell clumping. HEK-293T (ATCC CRL-3216) were cultured in standard DMEM (ThermoFisher) with 10% FBS. All cell lines were cultured at 37 °C at 5% CO_2_, seeded for approximately 50% confluency at the time of experiment, and were confirmed free of *Mycoplasma* contamination by the MycoAlert PLUS Mycoplasma Detection Kit (Lonza, Switzerland). Lentivirus was produced in 293T cells by transfection of dR8.91, pMD2.G, and the appropriate pLKO-derived vector (at ratios of 8 µg, 1 µg, and 8 µg, respectively, per 15 cm dish) with Trans-LT1 (Mirus Bio, Madison WI), according to the manufacturer’s instructions. Viral harvest media was supplemented with Viralboost (Alstem, Richmond CA), collected 2-3 days after transfection, and filtered through 0.44 µm polyvinylidene difluoride filters and either frozen for storage at −80 °C or used immediately for transduction.

### Generation of CRISPRi Cell Lines

All cell lines were transduced using virus-containing supernatant in the presence of 8 µg/ml polybrene (Millipore-Sigma, St. Louis MO). HepG2 expressing dCas9-KRAB were derived by transduction with lentivirus harboring SFFV-dCas9-BFP-KRAB, followed by two rounds of FACS for BFP-positive cells on a BD FACSAria II. dCas9-KRAB HepG2 with individual targeting sgRNAs were derived by transduction with lentivirus harboring the desired sgRNA, followed by 48 hrs of puromycin selection (2 µg/ml, InvivoGen, San Diego CA), prior to experiments.

### Quantitative Real-Time PCR

dCas9-KRAB HepG2 stably expressing an appropriate sgRNA were harvested, lysed, and total RNA was extracted via the RNeasy Mini Kit (Qiagen, Germantown MD). RNA was converted into cDNA using qScript cDNA SuperMix (QuantaBio, Beverly MA) following the manufacturer’s instructions. RT-qPCR was performed against indicated targets with PrimeTime qPCR primers (IDT, Coralville IA) using the SYBR Select Master Mix (ThermoFisher) according to the manufacturer’s instructions on a CFX96 Touch Real-Time PCR Detection System (BioRad, Hercules CA). Fold changes were calculated using ΔΔCt analysis, normalizing each sample to *B2M* controls, using CFX Maestro software (BioRad).

### Receptor Abundance Analysis

1-2 days prior to analysis, dCas9-HepG2 cells and derivatives were cultured in low-glucose DMEM with 5% lipoprotein deficient serum (Kalen Biomedical, Germantown MD). Prior to analysis, cells were dissociated with Accutase (Innovative Cell Technologies, San Diego CA), collected, washed in PBS (ThermoFisher), live-dead stained with Ghost Dye Red 780 (1:1000 dilution, Tonbo Biosciences, San Diego CA), washed, and then stained with the indicated antibody in FACS buffer (PBS with 1% FBS, 10 U/ml DNAse I, GoldBio, St. Louis MO) for 30 minutes on ice with gentle agitation. Cells were washed, resuspended in FACS buffer, filtered through 90 µm mesh (Elko Filtering, Miami FL) to give a single cell suspension, and placed on ice. Cells were then analyzed on either a BD Fortessa, BD LSRII, BD FACSAria II, or Beckman Coulter CytoFLEX, or sorted on a BD FACSAria II, depending on the experiment. In general, gating excluded cells positive for live-dead staining and included only the cells positive for the level of BFP expression induced by the CRISPRi/a v2 vector. FACS analysis and figure preparation was performed with FlowJo v10 (BD, Ashland OR).

### Genome-Wide CRISPRi Screen

The screen was conducted similarly to prior descriptions (*13–15*). Approximately 200 × 10^6^ dCas9-KRAB HepG2 were transduced with hCRISPRi-v2 top 5 sgRNAs/gene lentivirus at an MOI of ∼0.5, and with polybrene at 8 µg/ml, on day 1. Cells were grown on 15-cm dishes, subdivided into four replicates immediately upon transduction (biological duplicate for each screen), and reseeded every 3-4 days as necessary to avoid overconfluence. Cells were selected with puromycin (2 mg/ml) from day 2 through day 6. On day 5, cells for the LDLR sort were placed in DMEM with lipoprotein depleted serum (5%). On day 7, approximately 50 × 10^6^ cells from 2 replicates were live-dead stained and stained for LDLR as described above, and then two-way sorted on a BD FACSAria II for the top and bottom 33% by LDLR abundance. Cells were spun down, washed in PBS and frozen at −80 °C. On day 8, the sort was repeated except in one replicate, cells were stained for TFR instead of LDLR and then sorted as per above. Genomic DNA was isolated using a NucleoSpin Blood DNA extraction kit (Macherey-Nagel, Bethlehem PA). The sgRNA-containing region was PCR-amplified with NEBNext Ultra II Q5 MasterMix (NEB), acrylamide gel-purified, and size-selected by SPRI beads (Beckman Coulter, Indianapolis IN), all as previously described, prior to sequencing on an Illumina HiSeq 4000.

### Screen Processing

Sequencing data were aligned to the top5 library, counted, and quantified using the ScreenProcessing pipeline (accessed from https://github.com/mhorlbeck/ScreenProcessing*(14)* 4/25/2019). Phenotypes and Mann-Whitney *P* values were determined as previously described (*14*), with the phenotypes defined as follows: ρ indicated the comparison in high-abundance vs. low abundance cells, τ indicated the comparison in high-abundance vs. unsorted cells, and γ indicated the comparison in low-abundance vs. unsorted cells. Counts from 4 guides were removed from the final analysis as there was evidence of contamination from individually cloned plasmids (PCSK9_+_55505255.23-P1P2, HMGCR_+_74633053.23-P1P2, TFRC_-_195808987.23-P1P2, ACO1_-_32384733.23-P1P2). A hit threshold of 7 (normalized phenotype z score × −log10(p-value) ≥ 7) (*13*) was used to identify hits from ρ, τ, and γ phenotypes, which were then compiled. Identical analysis of the TFR screen was used to prioritize hits unique to LDLR regulation. Gene ontology analysis was performed using the PANTHER Classification System database (v15) (*28, 94*). For relaxation of the hit threshold for comparison to GWAS studies, a score of 6 was used. Cellular localization of hits was imputed by manual curation from UniProt (*95*) and the Human Protein Atlas (*55*).

### Human Genomic Analysis

Protein coding variants for hits validated at the individual sgRNA level were assayed in the UK Biobank (*96*) for associations with LDL-C. In the setting of a statin medication, LDL-C was divided by 0.7 as before (*30*). Genotyping and imputation was performed in the UK Biobank as previously described (*32*), and nonsynonymous protein coding variants with minor allele frequencies greater than 0.001 were considered. Efficient linear mixed models adjusting for age, sex, genotyping array, and principal components of ancestry were employed, using BOLT-LMM (*97*). Statistical significance was assigned at α = 0.05/117 = 0.000427 to account for multiple hypothesis testing.

### Validation Experiments of Individual sgRNAs

Cloning of protospacers, as described above, was performed in 96-well plate format until selecting individual colonies. Lentiviral production in 293T, transduction of dCas9-KRAB HepG2 with lentiviral sgRNA vectors, and receptor abundance and LDL uptake assays were similarly performed in 96-well plate format to maximize throughput.

### LDL Uptake Assays

Assays were performed as previously described(*33*) with the following modifications. dCas9-HepG2 cells harboring individual sgRNAs were treated similarly to receptor abundance analysis, except that prior to harvest, cells were washed and then treated with 5 µg/ml 1,1’-dioctadecyl-3,3,3’,3’-tetramethylindocarbocyanine perchlorate (DiI) labeled LDL (Kalen Biomedical) in low-glucose DMEM with 0.5% BSA (MilliporeSigma) for 1 hr at 37 °C. Cells were then washed, collected, and prepared for FACS analysis, as described above, but without antibody labelling.

### Pharmacologic Synergy Experiments

Receptor and LDL uptake assays were performed as described, with cells treated overnight with either simvastatin (6 µM, MilliporeSigma), PF-6846 (10 µM, MilliporeSigma), or DMSO vehicle (final concentration of 0.5%) overnight prior to analysis. Synergy scores were calculated by the following equation:

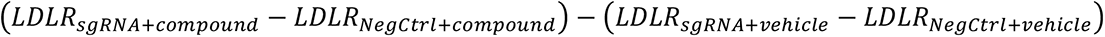

with *LDLR* obtained as the mean fluorescence, background subtracted from an unstained control and subsequently normalized to *LDLR_negctrl+vehicle_*. within a given experiment.

### Overexpression Experiments

HepG2 or engineered dCas9-HepG2 cell lines were seeded into 96 well plates at 5 × 10^4^ cells per well in HepG2 growth medium. After 24 hrs, cells were washed and changed into low-glucose DMEM with 5% lipoprotein-deficient serum. Each well was transfected with 100 ng of the appropriate CSDE1 overexpression construct, or vector control, in a total of 10 µL OptiMEM (ThermoFisher) using Lipofectamine 3000 (ThermoFisher) according to the manufacturer’s instructions. Cells were incubated at 37 °C with 5% CO_2_ for 72 hrs, and then harvested for LDL receptor expression analysis as above.

### mRNA Decay Experiments

Engineered dCas9-HepG2 cell lines harboring sgRNAs against CSDE1 (CSDE1_+_115300577.23-P1P2) or a negative control (Unassigned=negZNF335_-_44601297.24-all) were seeded into 12 well plates at 5 × 10^5^ cells per well in HepG2 growth medium. After 24 hrs, cells were washed and changed into sterol-depleting media (low-glucose DMEM with 5% lipoprotein-deficient serum) supplemented with 6 µM simvastatin. After an additional 24 hrs, actinomycin D (MilliporeSigma) was added at 5 µg/ml, and cells were harvested as described at the indicated timepoints.

### Immunoblots

Engineered dCas9-HepG2 cell lines harboring appropriate sgRNAs were grown in growth medium and harvested with 0.25% trypsin digestion. Cells were washed and lysed in lysis buffer on ice (50 mM Tris-HCl pH 7.4, 150 mM NaCl, 0.1% NP-40). Lysates were clarified at 21,000 × *g* for 10 min, and the supernatant was recovered. Equivalent amounts of lysates, as measured by BCA assay (ThermoFisher), were resolved on 4-12% Bis-Tris NuPAGE gels (ThermoFisher), transferred to nitrocellulose, probed with primary and secondary antibodies as noted (see Table) in 5% BSA in TBS-T, and visualized on an Odyssey imaging system (LI-COR, Lincoln NE).

### Dual-Luciferase Assays

Engineered dCas9-HepG2 cells were seeded into opaque white 96 well plates, at 2.2 × 10^4^ cells per well, in 100 µL growth medium the day prior to transfection. On day of transfection, medium was replaced or changed to sterol-depleted medium (low-glucose DMEM with 5% lipoprotein-deficient serum) with or without 6 µM simvastatin as appropriate. Each well was transfected with 100 ng of Luc2-Prom_LDLR_ based construct and 1 ng of secreted nanoluciferase control construct (pSS-NLuc) in a total of 10 µL OptiMEM using Lipofectamine 3000 according to the manufacturer’s instructions. 6 replicates were transfected per construct per experiment. After 48 hours at 37°C and 5% CO_2_, 10 µL of medium was removed from each plate and aliquoted into a separate 384 well plate. Firefly luciferase activity was evaluated in the plates containing the cells by adding an equal volume of a 2× firefly lytic assay buffer (100 mM Tris-HCl pH 7.7, 50 mM NaCl, 2 mM MgCl_2_, 0.17% Triton X-100, 10 mM DTT, 0.4 mM coenzyme A, 0.3 mM ATP, and 0.28 mg/ml luciferin (Goldbio)) (*98*). Nanoluciferase activity was evaluated from the conditioned medium using a non-lytic 2× coelenterazine (Goldbio) reagent as previously described (*91*). Raw luminescence was obtained on a SPARK plate reader (Tecan, San Jose CA) with 1 second integration time. Readout of firefly luciferase in each well was normalized to the corresponding secreted nanoluciferase control and data were visually inspected and cleaned to remove values from poorly transfected wells (formally defined by ROUT = 1%) during analysis.

### Zebrafish Handling, Maintenance, and Cas9-Ribonucleoprotein Knockdowns

All zebrafish studies were performed as previously described(*65*) with minor modifications. Briefly, wild type zebrafish embryos were injected at the one-cell stage with Cas9-RNP complexes and raised at 28 °C. Cas9-RNP complexes were prepared as previously described(*65*) using custom oligonucleotides against the indicated targets (Elim Biopharmaceuticals). Targeting of tyrosinase, which results in larval albinism, was used as an injection control. Larvae were fed a diet of Golden Pearls (5-50 micron, Brine Shrimp Direct, Ogden UT) 3× daily from 4 days post fertilization (dpf), fasted on 7 dpf to clear intestinal cholesterol, and harvested at 8 dpf. Larvae were collected, extensively washed, anesthetized in tricaine, and collected in groups of 10 per sample prior to storage at −80 °C. All zebrafish experiments were performed in accordance with IACUC-approved protocols at the University of California, San Francisco.

### Cholesterol Analysis of Zebrafish Homogenates

Total cholesterol levels were analyzed as previously described (*66*) with minor modifications. Briefly, frozen larvae were homogenized in PBS with a plastic pestle, and then clarified at 18,000 × *g* for 15 min. Supernatants were recovered and total protein content was analyzed by BCA assay. Homogenates were then analyzed, in duplicate, at the appropriate dilution (typically 1:12 in PBS) for total cholesterol content using a commercial fluorometric assay (Cayman Chemical, Ann Arbor MI). Fluorescence outputs were measured on a Tecan SPARK plate reader, and cholesterol concentrations were interpolated from a regression line calculated from a standard curve. Cholesterol levels were normalized to total protein content for analysis and subsequently to the scramble control for comparison between experiments.

### Mouse Handling, Maintenance, and shRNA Knockdowns

All mouse manipulations were performed in accordance with IACUC approved protocols at the University of California, San Francisco following guidelines described in the US National Institutes of Health Guide for the Care and Use of Laboratory Animals. 8-10 week old male C57BL/6 mice (The Jackson Laboratory, Bar Harbor ME) were placed on an atherogenic diet (1.25% cholesterol, 15% fat, 0.5% cholate, D12336i, Research Diets, New Brunswick NJ)(*68*) at the beginning of the experiment (week 0). At week 2, AAV8-packaged shRNA against mouse *Csde1* (NM_144901), *Pcsk9* (NM_153565), or scramble control (Vector Biolabs, Malvern PA) were diluted in sterile PBS to a concentration of 2 × 10^11^ genomes/ml. 100 µL of diluted AAV8 (2 × 10^10^ genomes/mouse) harboring the appropriate shRNA was administered to each mouse via tail vein injection. At week 4, and again at week 6, mice were fasted overnight and then underwent blood sampling via submandibular vein puncture. Approximately 50 µL of blood was collected into an EDTA-coated tube, centrifuged at 2000 × g for 10 min at 4 °C, and the plasma recovered and stored at −20 °C until further analysis. Total cholesterol of the plasma, after approximately 1:200 to 1:400 dilution in assay buffer, was evaluated by commercial fluorometric cholesterol assay (Cayman) as described above. At week 8, mice from the same exposure arm were re-dosed with AAV8 targeting either *Csde1* or scramble control. At week 10, the mice were again fasted overnight and then euthanized after CO_2_ narcosis followed by cervical dislocation. The abdominal cavity was opened with a ventral midline incision, the IVC was cannulated, and plasma was collected as described above. The liver and vasculature were perfused with PBS, and the samples of the liver were harvested. Tissue samples for RNA evaluation were placed in TRIzol (ThermoFisher) and those for protein analysis were flash frozen in liquid N_2_ and stored at −80 °C.

### Lipoprotein Fractionation

Plasma samples were thawed and centrifuged at 2000 × g for 10 min at 4 °C, and the supernatant recovered. 100 µL of individual mouse plasma was loaded onto a Superose 6 Increase 10/300 GL column (Cytiva, Marlborough MA) and eluted with PBS with 1 mM EDTA at 0.5 ml/min on an AKTA Pure chromatography system (Cytiva). Fixed 0.5 ml fractions were collected from 0.2 to 1 column volumes along the isocratic elution. Fractions were subjected to total cholesterol analysis and immunoblots as described above.

### RNA-seq Library Preparation

Total RNA was extracted from frozen liver samples using the Qiagen RNeasy Plus Universal mini kit followed by Manufacturer’s instructions (Qiagen). RNA samples were quantified using Qubit 2.0 Fluorometer (ThermoFisher) and RNA integrity was checked using Agilent TapeStation 4200 (Agilent Technologies, Palo Alto CA). Purified RNA was used for mouse qPCR experiments as described above. RNA sequencing libraries were prepared via polyA selection using the NEBNext Ultra RNA Library Prep Kit for Illumina using manufacturer’s instructions (NEB). Briefly, mRNAs were initially enriched with Oligod(T) beads. Enriched mRNAs were fragmented for 15 minutes at 94 °C. First strand and second strand cDNA were subsequently synthesized. cDNA fragments were end repaired and adenylated at 3’ends, and universal adapters were ligated to cDNA fragments, followed by index addition and library enrichment by PCR with limited cycles. The sequencing library was validated on the Agilent TapeStation (Agilent) and quantified by using Qubit 2.0 Fluorometer (Invitrogen) as well as by quantitative PCR (KAPA Biosystems, Wilmington, MA, USA). The sequencing libraries were clustered on a single lane of a flowcell. After clustering, the flowcell was loaded on the Illumina HiSeq instrument (4000 or equivalent) according to manufacturer’s instructions. The samples were sequenced using a 2×150bp Paired End (PE) configuration. Image analysis and base calling were conducted by the HiSeq Control Software (HCS). Raw sequence data (.bcl files) generated from Illumina HiSeq was converted into fastq files and de-multiplexed using Illumina’s bcl2fastq 2.17 software. One mismatch was allowed for index sequence identification. RNA library preparation and sequencing were conducted by GENEWIZ, LLC (South Plainfield, NJ).

### RNA-seq Analysis

All raw sequencing data underwent quality control checks with FastQC (v 0.11.8). Reads were mapped to the mm10 mouse reference genome using Rsubread (v 2.4.3) and assigned to Ensembl gene IDs. Ensemble gene IDs were then mapped to gene symbols using AnnotationDBI (v1.52.0). Gene expression was quantified using raw counts and differential expression gene testing was performed on the scramble-shRNA samples comparing the groups (n=3 in each group) at the highest and lowest levels of raw eGFP expression with EdgeR (*99, 100*) (v.3.32.1) using the glmQLFit method, default settings (*101*). Statistical significance was set at 5% false discovery rate (FDR; Benjamini-Hochberg). Differential expression gene testing was then performed on the *Csde1*-shRNA and scramble-shRNA at the highest levels of eGFP expression with the overlap of differentially expressed genes identified between these two analyses subsequently removed. Functional enrichment gene-set analysis for GO (Gene Ontology) terms was performed using Enrichr (*102*) via the enrichR R package (v.3.0). Heatmaps were generated using the Bioconductor package ComplexHeatmap (*103*) (v.2.6.2) using log2-transformed CPM values (counts-per-million; values shown are log2-transformed and row-normalized). Volcano plots were generated using the Bioconductor package EnhancedVolcano (v.1.2.0).

### Statistical Analysis

Fluorescence values from gated populations in flow cytometry experiments were background corrected by unstained controls and were normalized to the values of the cell line harboring negative control sgRNA. Normalized data were then grouped by the Cochrane method (*104*), and values for cell lines transduced with individual sgRNAs were compared those of the negative control by T-test with Holm-Sidak correction. For comparison of one-phase decay regression curves in mRNA decay experiments, the extra sum-of-squares F test was used. Pairwise testing to controls was performed in all other experiments using Welch’s T-test with Holm-Sidak correction unless otherwise noted. One or two-way ANOVA with Tukey’s or Sidak’s multiple comparisons tests was used when comparing across all groups in a particular experiment. Adjusted p values < 0.05 were considered significant. Statistical analysis was performed using Prism 7 (GraphPad Software, San Diego CA). All experiments were replicated thrice unless otherwise noted.

## Key Resources Tables

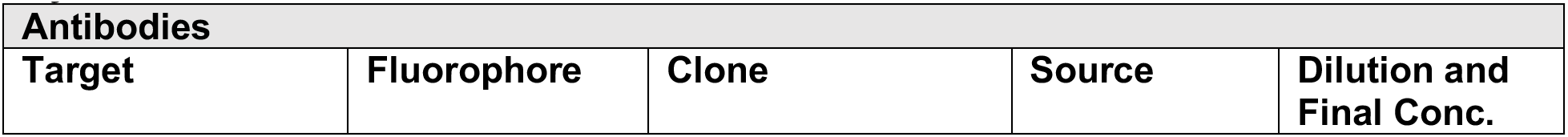
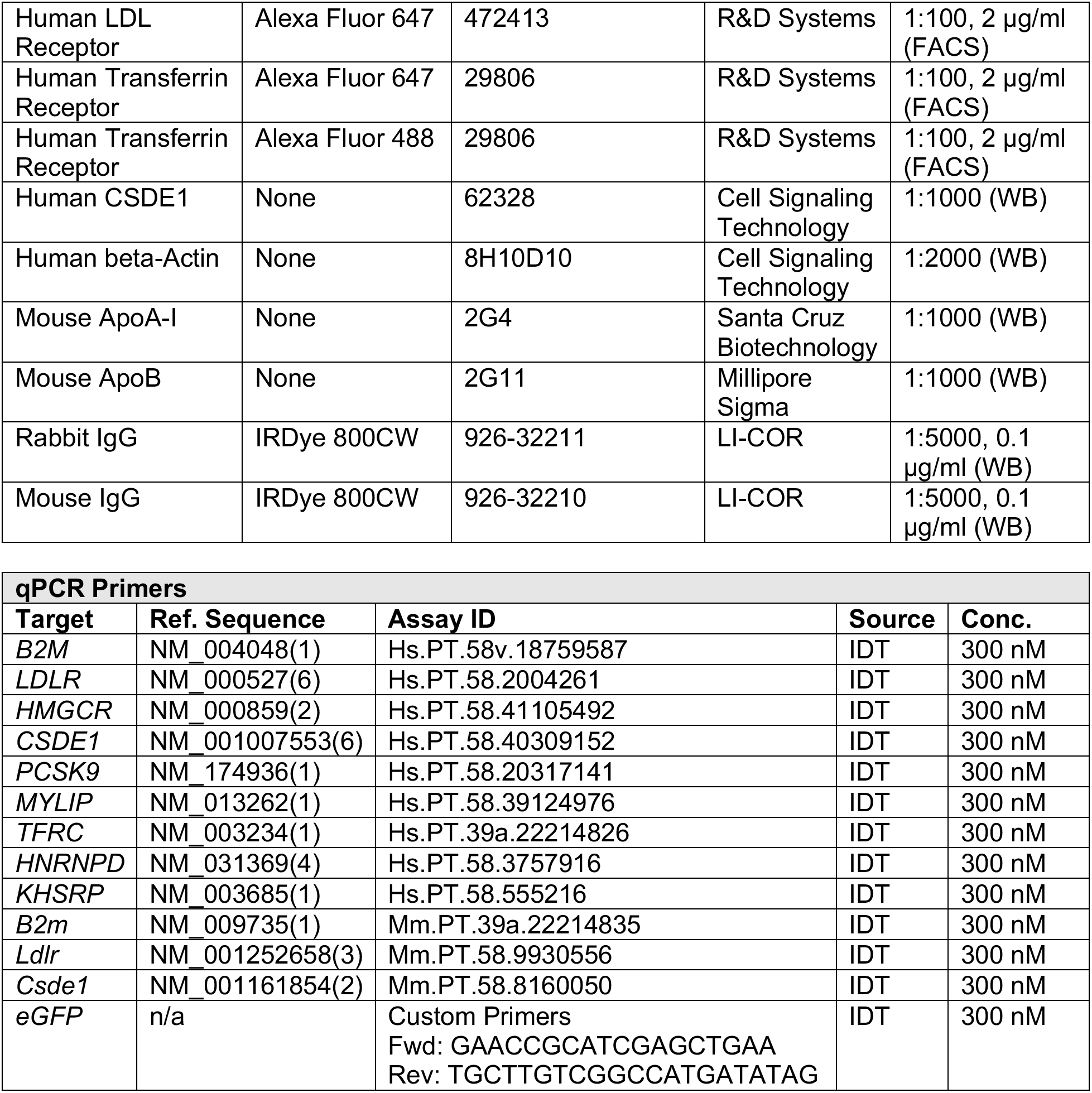

## Acknowledgements

We thank Max Horlbeck for guidance with the CRISPRi system, Jonathan D. Brown for helpful discussion on *in vivo* experimental designs, and Paul Cheng and Richard Baylis for helpful discussion on histologic insights and *in vivo* experimental planning. Plasmids for the CRISPRi system were a generous gift from Luke Gilbert and Jonathan Weissman. We thank the Gladstone Institutes Flow Cytometry core facility for their assistance with flow cytometry experiments. UK Biobank analyses were conducted using the UK Biobank resource under application 7089.

## Author Contributions

Overall study design: J.S.C. Execution of *in vitro* screen and data processing: G.A.S. Genomic analyses: Ak.P., P.N. *In vitro* validation and synergy experiments, and analysis of mouse blood samples: J.S.C. Execution of zebrafish gene knockdowns: B.H.L., R.S.W. Oversight of zebrafish husbandry: R.S.W., B.L.B. Planning and execution of *in vivo* mouse experiments: Ar.P. Mouse handling, blood, and tissue collection: Ar.P., Y.C.L., T.N., N.S. Processing of RNA-seq data: Ar.P., An.P. Analysis of mouse samples: L.L., R.J. Critical data review and analysis: B.L.B, D.S., K.M.S. Preparation of manuscript: J.S.C. Critical review and revision of manuscript: All authors.

## Resource and Data Availability Statement

Additional supporting data are available upon request from the corresponding author. All requests for raw and analyzed data, and materials, including plasmids or cell lines, generated in this study will be responded to promptly. UK Biobank data is available by application to the UK Biobank.

## Supplementary Materials

**Supplementary Figure 1:**
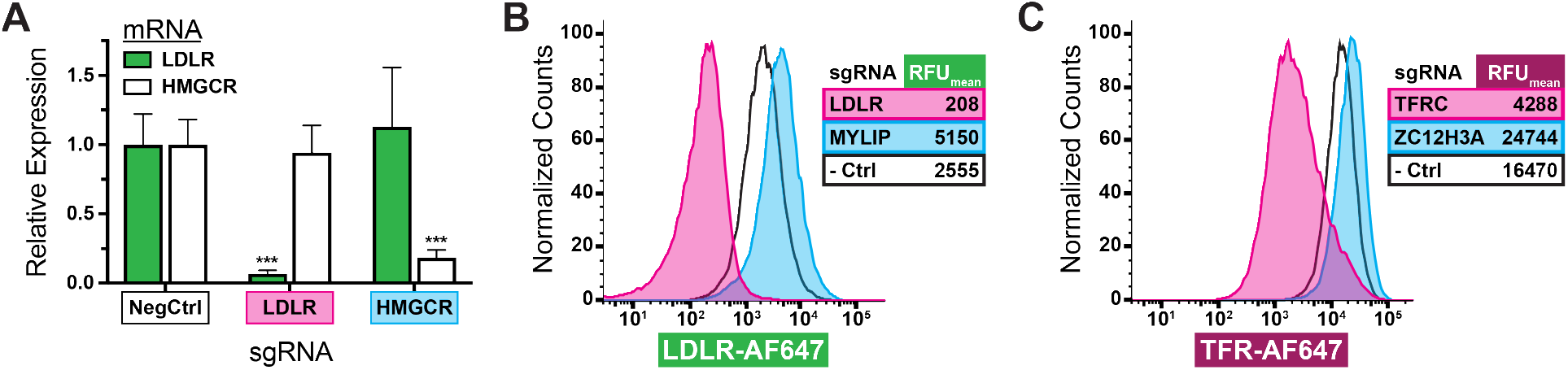
Validation of dCas9-KRAB-HepG2 Cells. *A)* Relative expression, by qPCR, of *LDLR* and *HMGCR* in engineered dCas9-KRAB HepG2 cells transduced with sgRNAs targeting the indicated genes. *B2M* used as qPCR control. Error bars indicate 95% confidence intervals. *** = p < 0.001 by Holm-Sidak corrected T-test, comparing to negative control sgRNA of the same target. *B)* Flow cytometry analysis, by surface labelling with anti-LDLR-AF647, of dCas9-KRAB HepG2 cells transduced with sgRNAs targeting the indicated genes. Mean fluorescence shown in inset. *MYLIP* (IDOL) is an E3 ligase which ubiquitinates the LDLR, leading to lysosomal degradation(*26*). *C)* Flow cytometry analysis as in *B* but transduced with indicated sgRNAs and labelled with anti-TFR-AF647. *ZC3H12A* (REG1) is an endoribonuclease that accelerates the degradation of TFR mRNA(*27*).

**Supplementary Figure 2:**
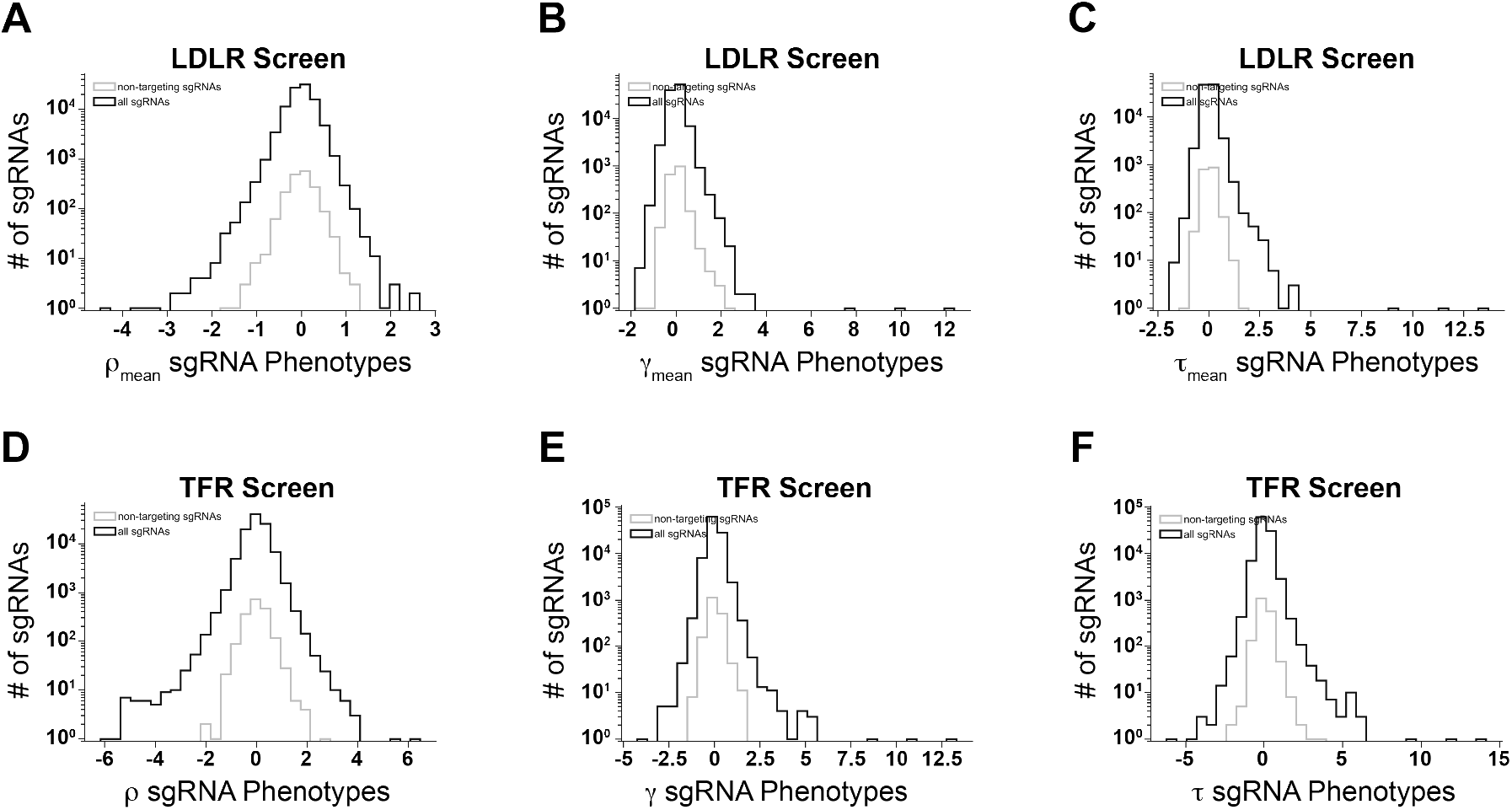
Recovered sgRNAs from Screening Phenotypes. Distribution of number of guide RNAs recovered by phenotype in both LDLR (*A-C*) and TFR (*D-F*) screens. ρ (*A,D*) indicates log_2_ fold enrichment for sgRNA in high receptor abundance cells compared to low receptor abundance cells. γ (*B,E*) indicates log_2_ enrichment for sgRNA in low receptor abundance cells compared to unsorted population. τ (*C,F*) indicates log_2_ enrichment for sgRNA in high receptor abundance cells compared to unsorted population. Mean results are reported for the 3 replicates of the LDLR screen (*A-C*).

**Supplementary Figure 3:**
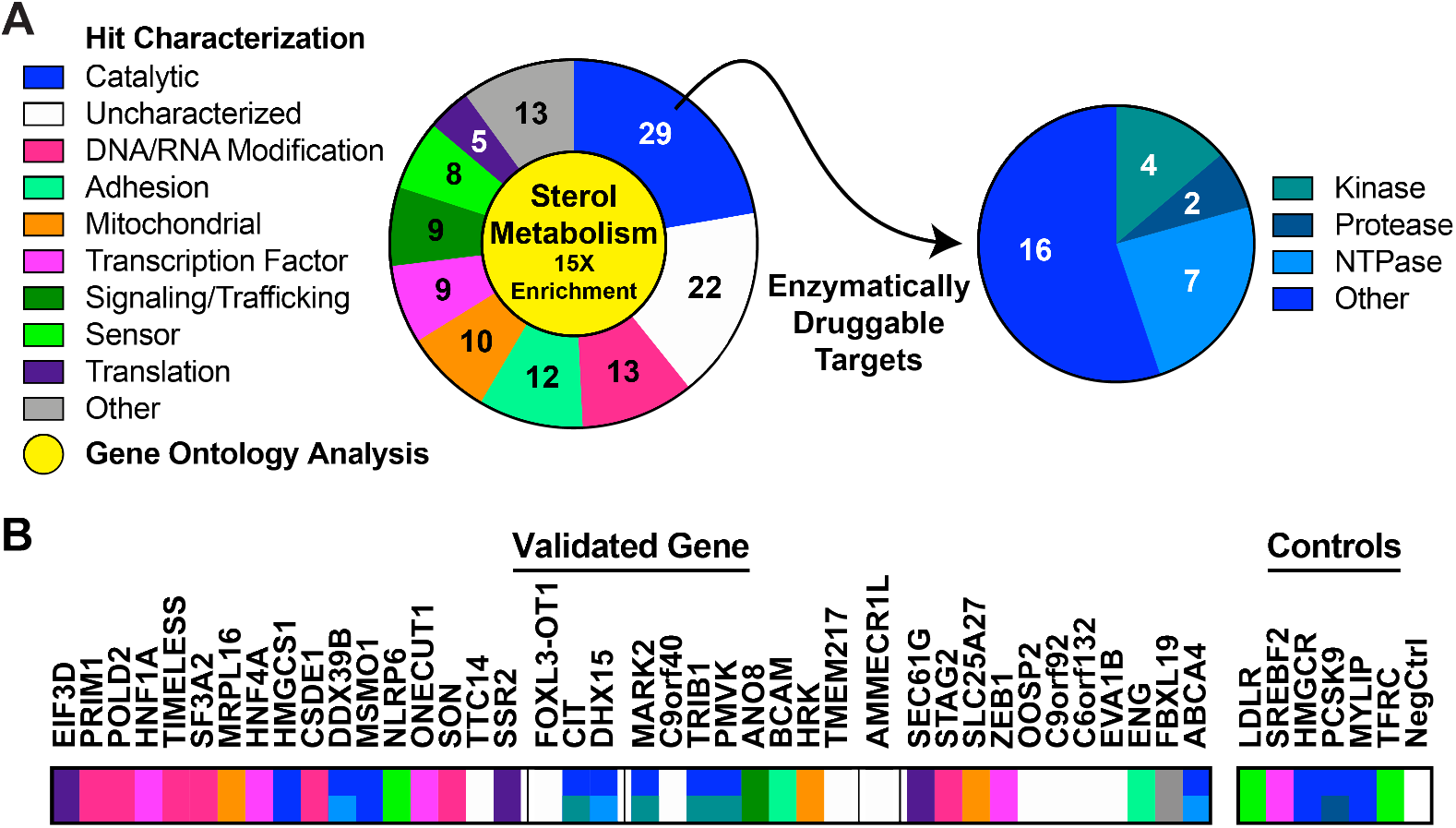
Gene Ontology and Localization Analysis. *A)* Characterization of hits from the LDLR screen based on gene ontology (GO) and localization, along with results from GO enrichment analysis (yellow center). Note that multiple genes fall into more than one category. *B)* Primary classification of the 40 LDLR hits independently validated outside of the pooled screen and displayed, as in Fig. 2, according to the color codes in *A*.

**Supplementary Figure 4:**
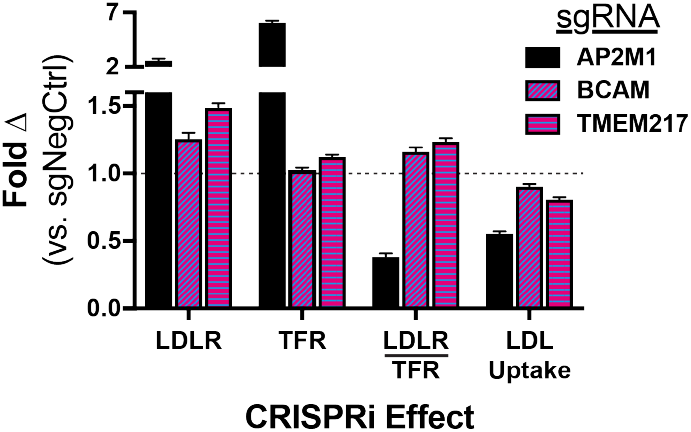
Selective LDLR Effect of Transmembrane Proteins. Flow cytometric readout of receptor abundance and LDLR function assays, using CRISPRi knockdowns against genes thought to be involved in endocytosis. Data, which represents 3 to 4 independent experiments, are normalized to readout of negative control sgRNA within each experiment. Error bars represent 95% confidence intervals. Note the discontinuous Y axis.

**Supplementary Figure 5:**
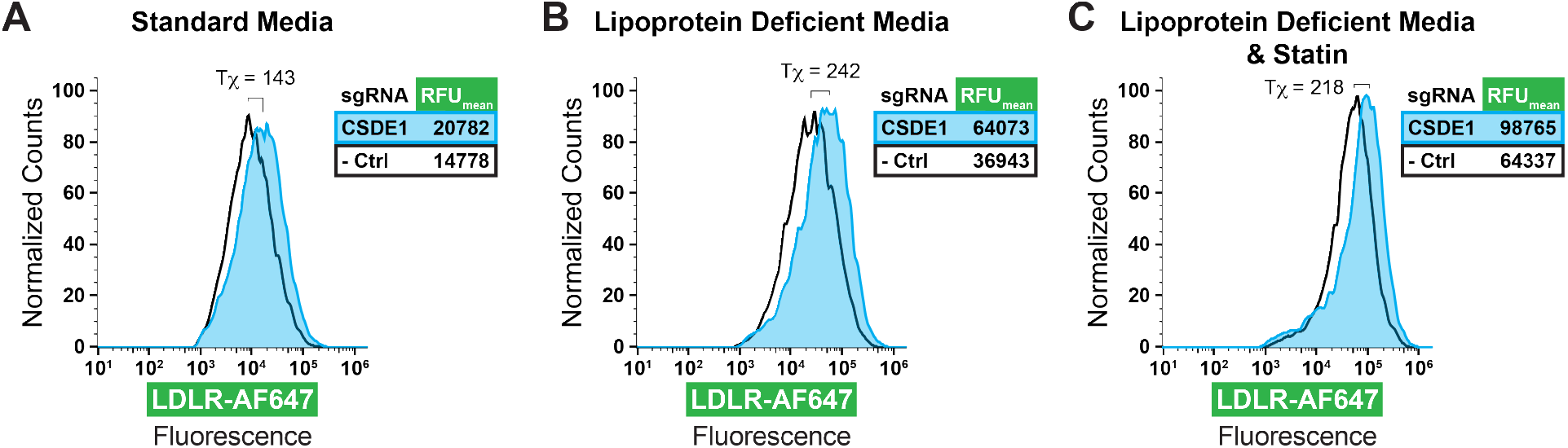
Effect of Sterol Conditions on CSDE1 Knockdown. Flow cytometry histograms showing AF647 conjugated anti-LDLR antibody labelling (as proxy for LDLR abundance) of engineered dCas9-KRAB HepG2 cells transduced with indicated sgRNAs and grown in standard growth media (*A*), lipoprotein-deficient media (*B*), or lipoprotein deficient media with a concomitant statin (*C*). Tχ metric (FlowJo v10)(*105–107*) shown on graph, and mean fluorescence shown in insets.

**Supplementary Figure 6:**
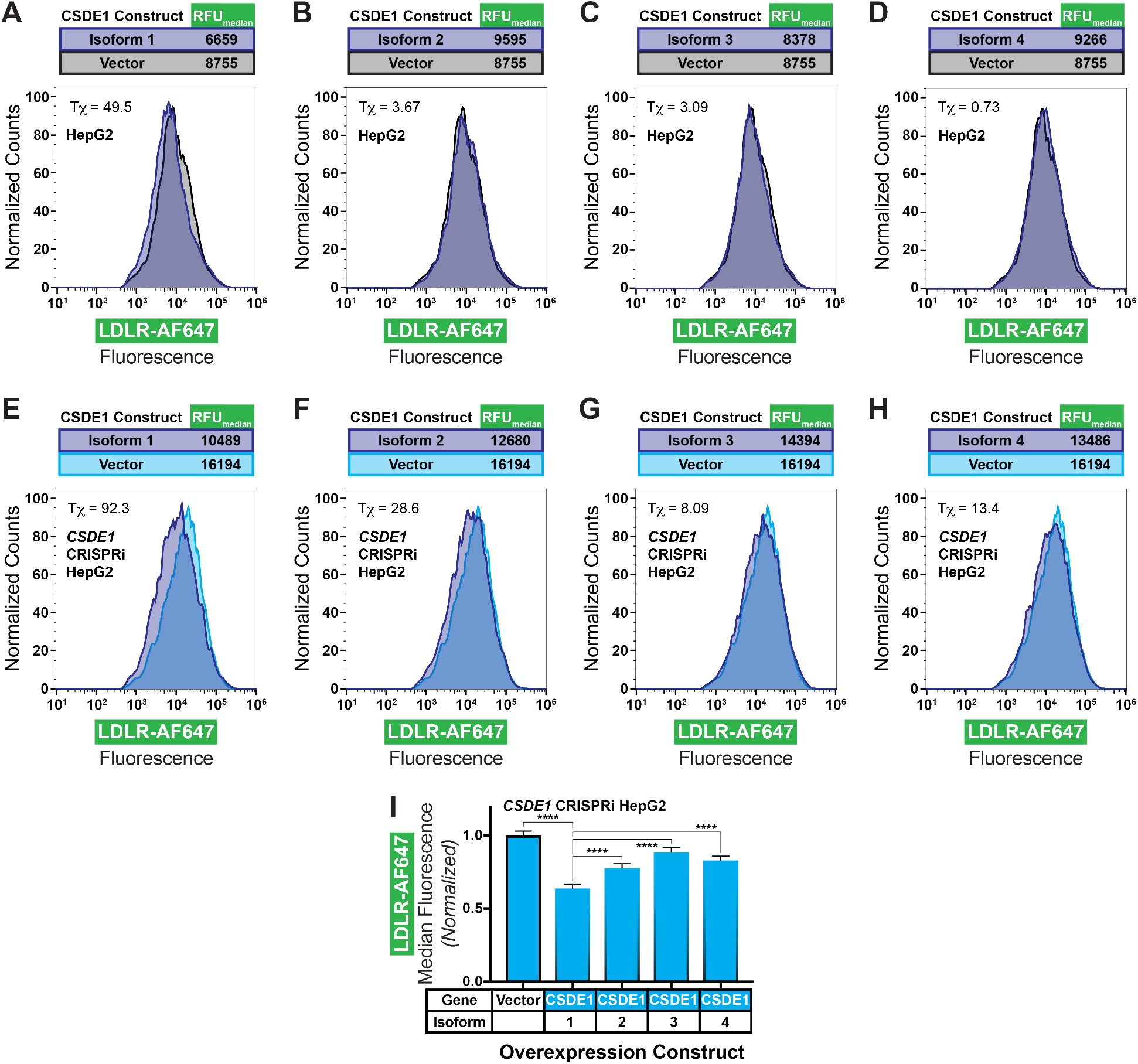
Effect of CSDE1 Overexpression. Flow cytometry histograms showing AF647 conjugated anti-LDLR antibody labelling (as proxy for LDLR abundance) of HepG2 cells (*A-D*) or engineered dCas9-KRAB HepG2 cells transduced with *CSDE1* targeting sgRNA (*E-H*) and transfected with an overexpression construct encoding the indicated CSDE1 isoform or vector alone. Τχ metric (FlowJo v10)(*105–107*) shown on graphs, and median fluorescence shown above. Quantified relative LDLR abundance of normalized median fluorescence of the data from *e-h* shown in *i*, with one-way ANOVA with Tukey’s multiple comparisons test. Error bars indicate 95% confidence intervals. **** = p < 0.0001.

**Supplementary Figure 7:**
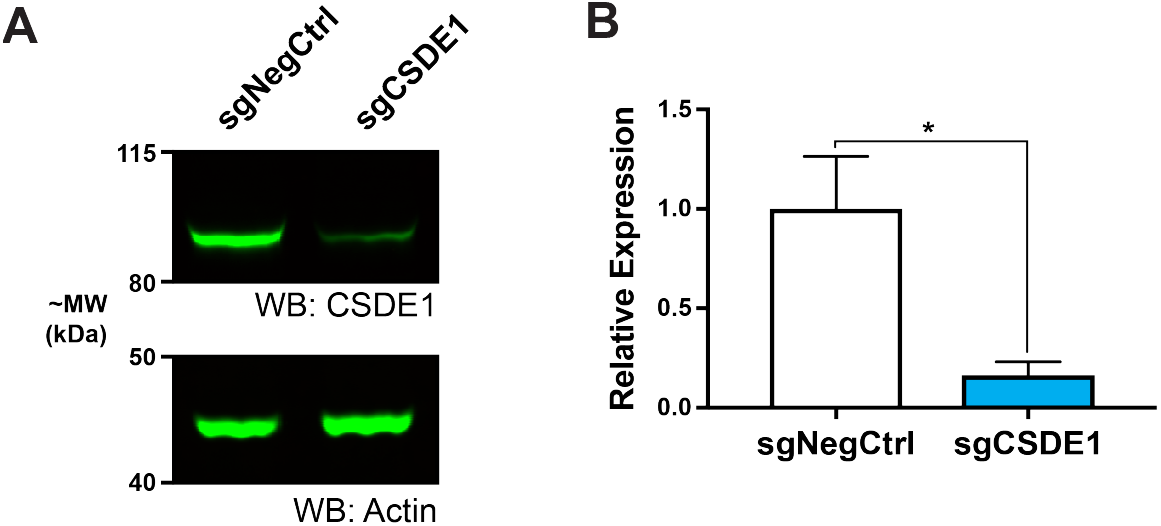
CSDE1 Knockdown at Protein Level. *A*) Representative immunoblots of lysates of dCas9-KRAB HepG2 cells harboring indicated guide RNA. CSDE1 shown above, and β-actin (loading control) shown below. *B*) Quantification of relative abundance of CSDE1 (normalized to loading control) shown in immunoblot in *A*. Data includes 3 independent experiments. * indicates p < 0.05 by Welch’s T-test.

**Supplementary Figure 8:**
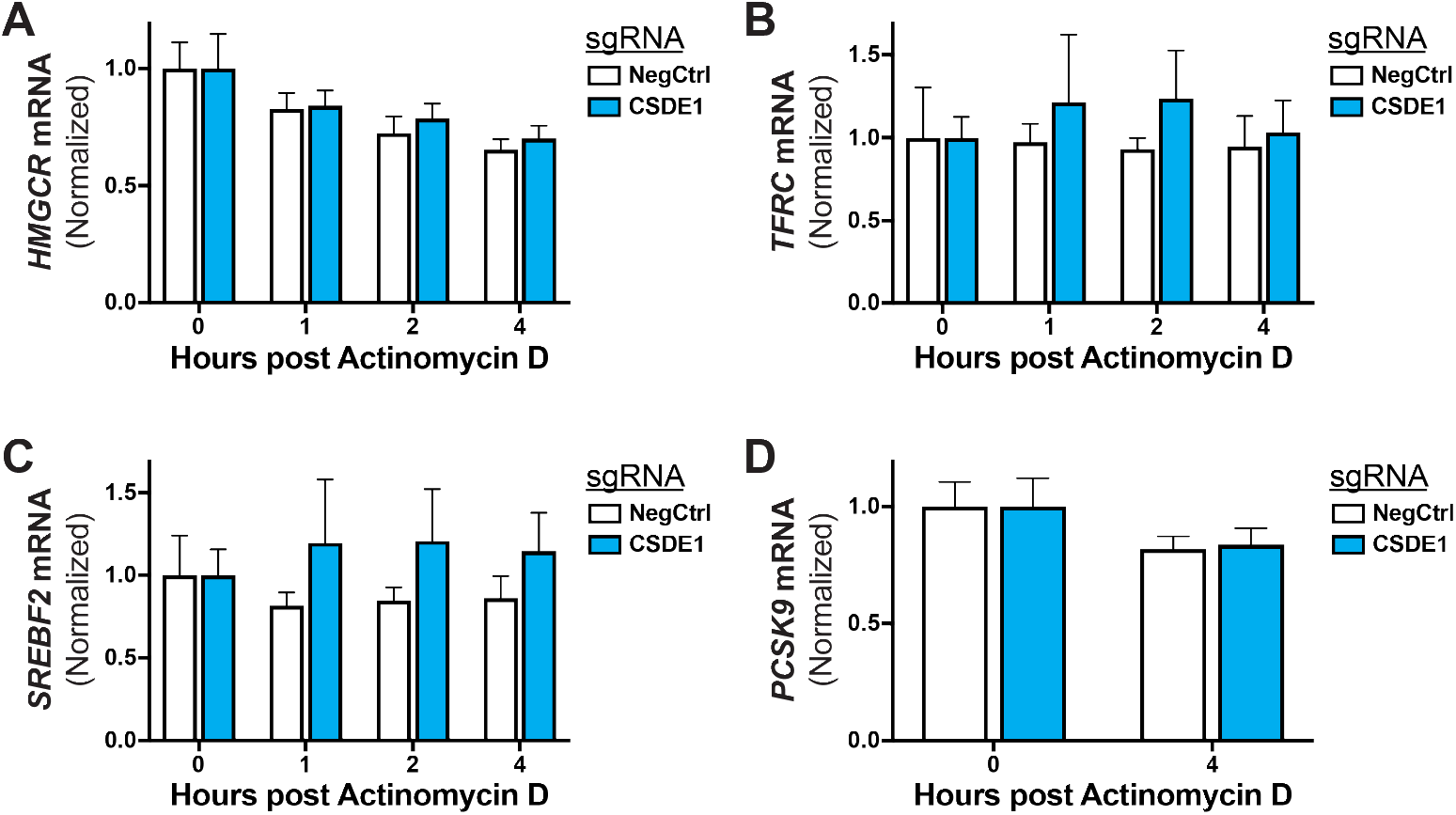
Effect of *CSDE1* Knockdown on Decay of Non-*LDLR* Transcripts. Relative expression, by qPCR, of *HMGCR (A), TFRC (B), SREBF2 (C), or PCSK9 (D)* mRNA in dCas9-KRAB HepG2 cells transduced with indicated sgRNAs and subjected to arrest of transcription with actinomycin D. Data are normalized to results at T=0 within the sgRNA evaluated to illustrate the change in time. Data represent summary results from 3 independent experiments. Error bars = 95% confidence intervals. All pairwise comparisons (unpaired t-tests with Holm-Sidak correction) are nonsignificant.

**Supplementary Figure 9:**
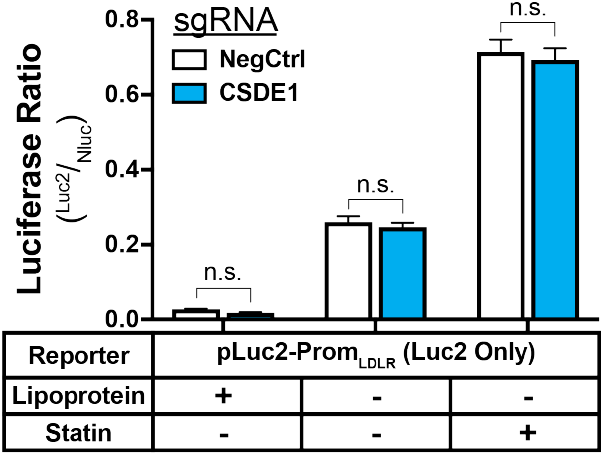
Physiologic Response of Luciferase Reporter System. Relative ratiometric luciferase activity of dCas9-KRAB HepG2 (CRISPRi) cells transiently transfected with unmodified Luc2 construct under the *LDLR* promoter (pLuc2-Prom_LDLR_, Fig. 4E) and secreted Nluc reporter under the CMV promoter and subjected to indicated media conditions. Data represent summary results from 3 independent experiments. n.s. = non-significant. Two-way ANOVA with Sidak’s multiple comparisons test was used.

**Supplementary Figure 10:**
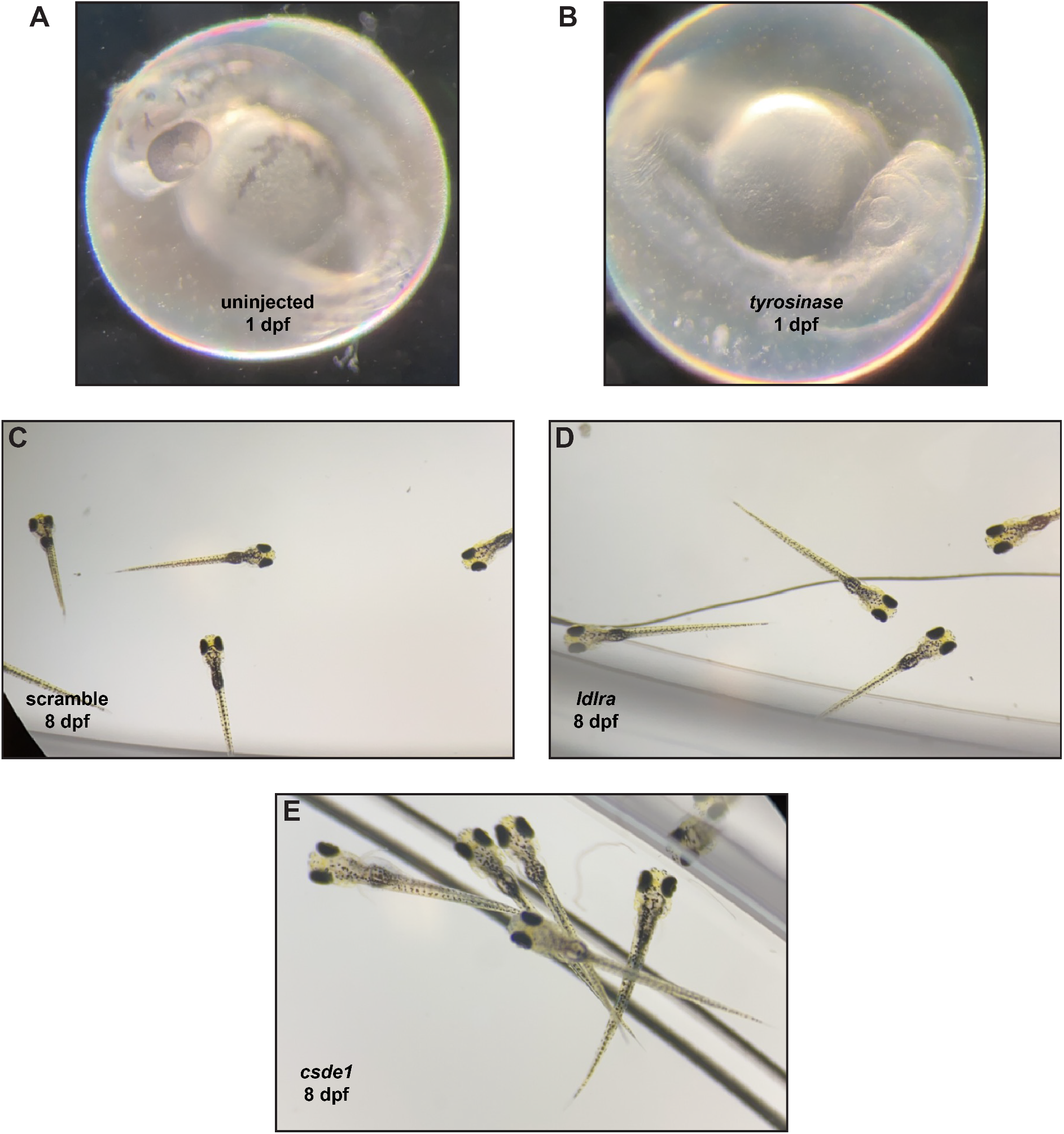
Visual Phenotypes of Zebrafish Cas9-sgRNA Saturation Gene Disruption. *A,B*) Representative microscopic images of zebrafish larvae at 1 day post fertilization without (*A*) or with (*B*) injected Cas9 and redundant guides against tyrosinase control performed concomitantly with each zebrafish experiment. Albinism is the readout for successful injections. *C-E)* Representative microscopic images of zebrafish larvae at 8 dpf and with injected Cas9 and guides against scramble controls (*C*), *ldlra* (*D*), and *csde1* (*E*).

**Supplementary Figure 11:**
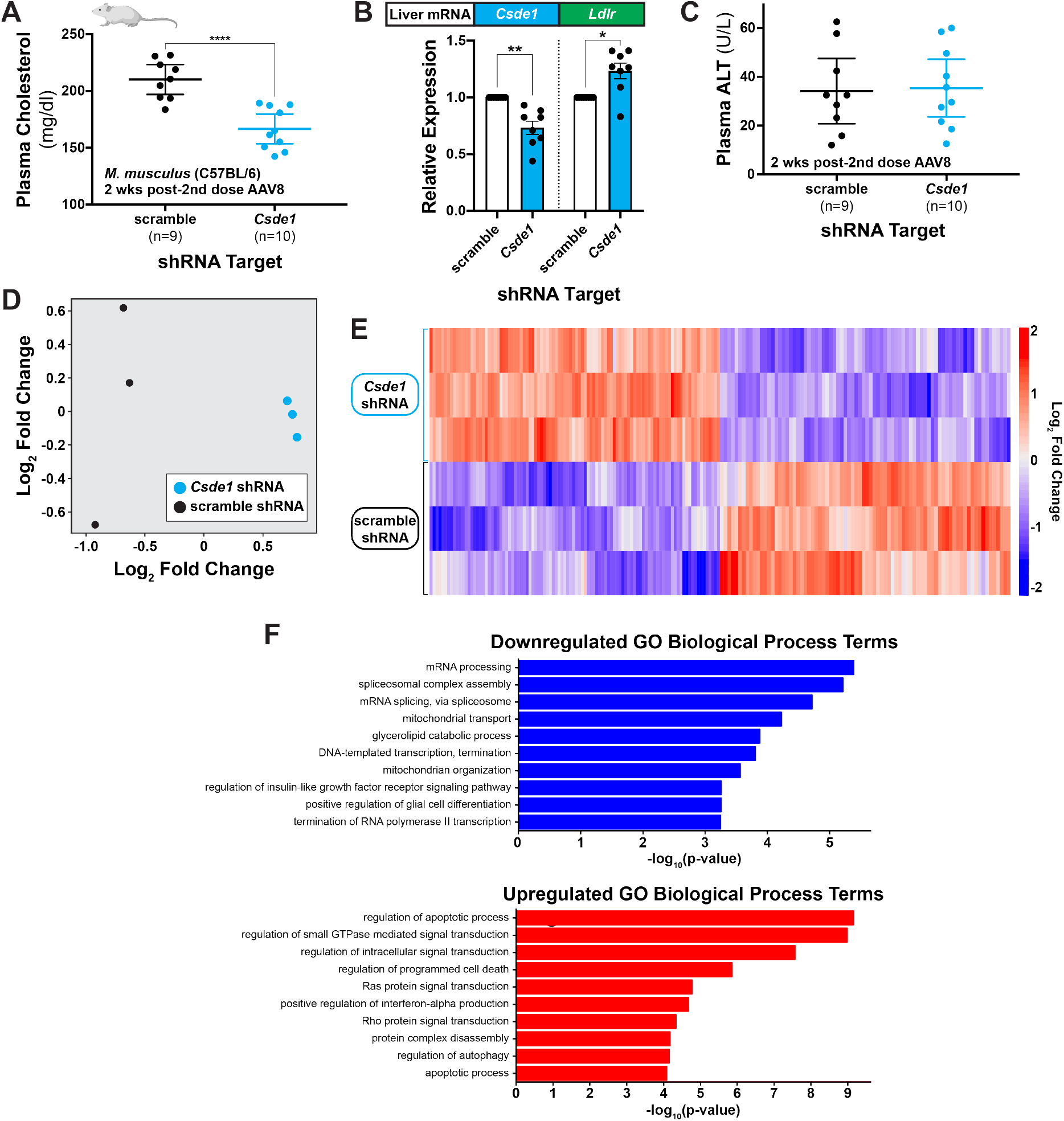
Effects of *in vivo Csde1* Disruption in Mice. *A)* Total fasting plasma cholesterol of C57BL/6 mice on an atherogenic diet, 2 weeks after transduction with second dose of AAV8-packaged shRNA against indicated target (8 weeks after first dose). Welch’s t-test shown. *B)* Relative expression, by qPCR, of indicated mRNA targets from mouse liver tissue from the indicated treatment arms. Expression normalized to *B2m* as the housekeeping control. Welch’s T-test shown. Error bars = standard error of the mean. *C)* Plasma alanine aminotransferase activity (ALT) of mice from *A*. Welch’s T-test revealed no significant difference between intervention arms. *D)* Unsupervised cluster analysis of individual mice analyzed for differential gene analysis. *E)* Heatmap of top 100 differentially expressed genes (by p value) of the individual mice analyzed by RNA-seq. Mice transduced with AAV8-scramble-shRNA at the bottom and those transduced with AAV8-*Csde1*-shRNA at the top. *F)* Top 10 biological process GO terms (by uncorrected p value) identified from all statistically significant differentially expressed genes downregulated (blue) or upregulated (red) by *Csde1* shRNA treatment. *All)* Each data point represents an individual mouse. * = p < 0.05, ** = p < 0.01, and **** = p < 0.001. Error bars = 95% confidence intervals unless noted otherwise.

Table S1: LDLR Screen Data by Gene *(provided as an Excel file)*

Table S2: LDLR Screen Data by Guide *(provided as an Excel file)*

Table S3: TFR Screen Data by Gene *(provided as an Excel file)*

Table S4: TFR Screen Data by Guide *(provided as an Excel file)*

Table S5: LDLR Screen Hits *(provided as an Excel file)*

Table S6: TFR Screen Hits (provided as an Excel file)

**Supplementary Table 7:** Baseline Characteristics of UK Biobank Participants in Genomic Association Analyses. Continuous values are presented as mean (standard deviation) except for triglycerides which is given as median (Q1-Q3) due to the skewness of the triglyceride distribution. Categorical data are presented as count (percentage). BMI = body-mass index; HDL = high-density lipoprotein; LDL = low-density lipoprotein.

| Metric | Value |
| --- | --- |
| Age (y) | 56.9 (7.9) |
| Sex | 179,963 (46.1%) |
| European ancestry | 376,358 (96.4%) |
| Cholesterol (mg/dl) |  |
| Total | 221.1 (44.3) |
| HDL | 56.1 (14.8) |
| LDL | 138.1 (33.7) |
| Triglycerides (mg/dl) | 132.6 [93.6-191.5] |
| Statin Rx | 64,004 (16.4%) |
| BMI (kg/m<sup>2</sup>) | 27.4 (4.8) |
| Systolic blood pressure (mmHg) | 140.2 (19.7) |
| Diastolic blood pressure (mmHg) | 82.3 (10.7) |
| Current smoker | 39,736 (10.2%) |
| Diabetes mellitus type 2 | 25,349 (6.5%) |
| Coronary artery disease | 18,204 (4.8%) |

Table S8: Validation Data by Guide *(provided as an Excel file)*

Table S9: Pharmacology Synergy Data by Guide *(provided as an Excel file)*

Table S10: Differentially Expressed Genes by *in vivo* RNA Seq *(provided as an Excel file)*

Table S11: Enriched GO Terms by *in vivo* RNA Seq *(provided as an Excel file)*

